# Single-cell characterization of 3D CLL spheroid model that recapitulates lymph node microenvironment-driven phenotypes and in vivo drug responses to ibrutinib

**DOI:** 10.64898/2026.10.31.763314

**Authors:** Marco V Haselager, Danial Lashgari, Aldo Jongejan, Jamie Lee Roos, Marjolein Dijkema, Julie Dubois, Fleur S Peters, Arnon P Kater, Perry D Moerland, Eric Eldering

**Author notes:** Shared first authorship (Marco V Haselager, Danial Lashgari). Corresponding author: Eric Eldering, Department of Experimental Immunology Amsterdam UMC, 1081BT, Amsterdam, the Netherlands, Fax: not available.

## Abstract

Lymph node (LN) microenvironmental signals promote chronic lymphocytic leukemia (CLL) cell survival, proliferation, and treatment resistance. We previously developed a 3D culture model that recapitulates key features of this protective LN niche. Using single-cell RNA sequencing of ex vivo paired peripheral blood and LN samples, together with matched patient-derived CLL spheroids, we investigated how microenvironmental cues shape CLL phenotypes and their response to ibrutinib. A total of 3 functional CLL states (resting, activated, proliferating) were identified, with expanded populations of activated and proliferating subsets in 3D cultures. After 5 days of in vitro ibrutinib treatment, CLL cells shifted toward a resting phenotype, reflecting published responses in patients at 6 months. Activated CLL cells showed relative resistance to ibrutinib, associated with TLR9-MALT1-MYC pathway activation. The 3D model recapitulated tissue-specific T cell phenotypes observed in patients, and computational analyses predicted several T-to-CLL cell-mediated interactions supporting the LN niche. Collectively, these data reveal distinct ibrutinib responses among CLL subsets and establish the 3D model as a physiologically relevant platform.

## Introduction

Beyond B cell receptor signaling, CLL cells depend on multiple microenvironmental signals that promote their proliferation and survival, mediated by interactions within the lymph node (LN) involving T cells, myeloid cells, stroma and soluble factors^1–3^. CLL cells continuously traffic between LNs and peripheral blood (PB), with enrichment of activated and proliferating cells in LN compartments, whereas the PB compartment predominantly contains resting CLL cells. This trafficking equilibrium is disrupted by therapeutic interventions that target microenvironmental interactions, most notably the BTK inhibitor ibrutinib.

Ibrutinib has shown remarkable clinical efficacy with high response rates, including in high-risk patients with genomic aberrations^4,5^. Its clinical success is attributed to disrupting B cell receptor signaling as well as the release of CLL cells from LNs into the PB and preventing migration back into the LNs. Due to the ibrutinib-induced loss of supportive LN cues and the lack of survival signals in the PB, CLL cells will eventually die^5–7^. Patients may experience relapse during ibrutinib treatment, suggesting that persisting tumor cells are protected at LN sites and may subsequently drive disease progression^8,9^.

Understanding what happens to LN-residing CLL cells under ibrutinib is crucial to explain why and how patients eventually relapse, but this is not easily studied in vivo due to material scarcity after treatment-induced redistribution of CLL cells into the PB. An arguably bigger drawback is that CLL is widely studied using PB samples and that many signals that drive CLL activation, proliferation and drug resistance are not recapitulated in standard in vitro models^3^.

We developed a 3D CLL culture model that utilizes PB-derived patient samples and incorporates CLL, T cells and myeloid cells in a spatially organized, LN-mimicking context^10^. This model results in expanded populations of activated and proliferating CLL cells, allowing for parallel investigation of proliferation and drug resistance. In the current study, we aim to dissect the transcriptional and functional responses of LN-resident as well as PB CLL cells to ibrutinib using single-cell RNA sequencing (scRNA-seq) of CLL spheroids and paired patient samples. First, we characterized CLL and T cell phenotypes and dissected tissue-specific transcriptional profiles. Second, we investigated the ibrutinib responses across CLL subpopulations within the 3D model. Third, we compared the ibrutinib response in the 3D CLL model to the in vivo response of matched patient samples.

In summary, we show that functional CLL state strongly influences transcriptional profiles and ibrutinib responses. We find that activated CLL cells showed reduced transcriptional responses to ibrutinib compared to resting and proliferating CLL cells, in which the TLR9-MALT1-MYC axis may represent a potential escape mechanism during BTK inhibition that warrants further investigation. Finally, we inferred T-CLL interactions associated with niche-dependent effects of ibrutinib.

## Material & Methods

### Patient material

With written informed consent of patients, paired PB and LN samples were collected in the context of the HOVON 158 clinical trial (ClinicalTrials.gov, NCT04639362), a fully recruited, open-label, non-randomized phase 2 trial at 17 hospitals in the Netherlands and Denmark^11^. In the current study, we used PB and LN samples collected at baseline as well as PB samples after 3 months of ibrutinib monotherapy. Blood mononuclear cells were obtained after Ficoll density gradient centrifugation (Pharmacia Biotech, Roosendaal, The Netherlands), were cryopreserved and stored as previously described^12^. In the case of LN samples, hollow needle biopsies were collected at baseline, immune cells were filtered through a cell strainer and similarly processed by Ficoll density gradient centrifugation and cryopreservation as previously described^12^. Cells were stained with the following antibodies: CD8, CD56, CD5, CD19, TCRγδ, CD3, CD4, and CD20, and measured by flow cytometry on a Canto II flow cytometer (BD Biosciences, Franklin Lakes, New Jersey [NJ]). All patient samples included in this study are listed in Supplemental Table 1.

### 3D culture

Peripheral blood mononuclear cells (PBMCs) and paired LN-derived samples of CLL patients were plated in ultra-low attachment (ULA) plates and centrifuged for 10 minutes at 1000 rpm and cultured for 5 days. Cells were cultured at 37°C and 5% CO2 using standard cell culture techniques. Cells were cultured in Iscove’s Modified Dulbecco’s Medium supplemented with 10% fetal calf serum and 1% penicillin/streptomycin. Cells were stimulated with a B cell stimulation cocktail consisting of 25 ng/mL interleukin (IL)-2, IL-15, and IL-21 including 1 µg/mL CpG in the absence or presence of 100 nM ibrutinib (SelleckChem, cat. no. S2680). For each culture condition, 24 spheroids were cultured and pooled.

### Single-cell RNA sequencing

Cells were sorted at a ratio of 60% CLL cells versus 40% non-CLL cells. Cell hashing with BioLegend TotalSeq antibodies (B0251, B0252, B0253, B0254, B0255, B0256, B0257) enabled multiplexing of 7 conditions per patient (Fig. 1A). Each multiplexed pool was split and loaded onto 2 wells of a 10x Genomics Chromium chip, targeting recovery of 16,000 single cells per well. Single-cell gene expression and hashtag libraries were generated with the Chromium Next GEM Single Cell 3’ Kit v3.1 and the Dual Index Kit NN Set A, respectively. Libraries were sequenced on a NovaSeq X Plus Instrument targeting 30,000 reads per cell for gene expression and 4,000 reads per cell for hashtags.

**Figure 1.**
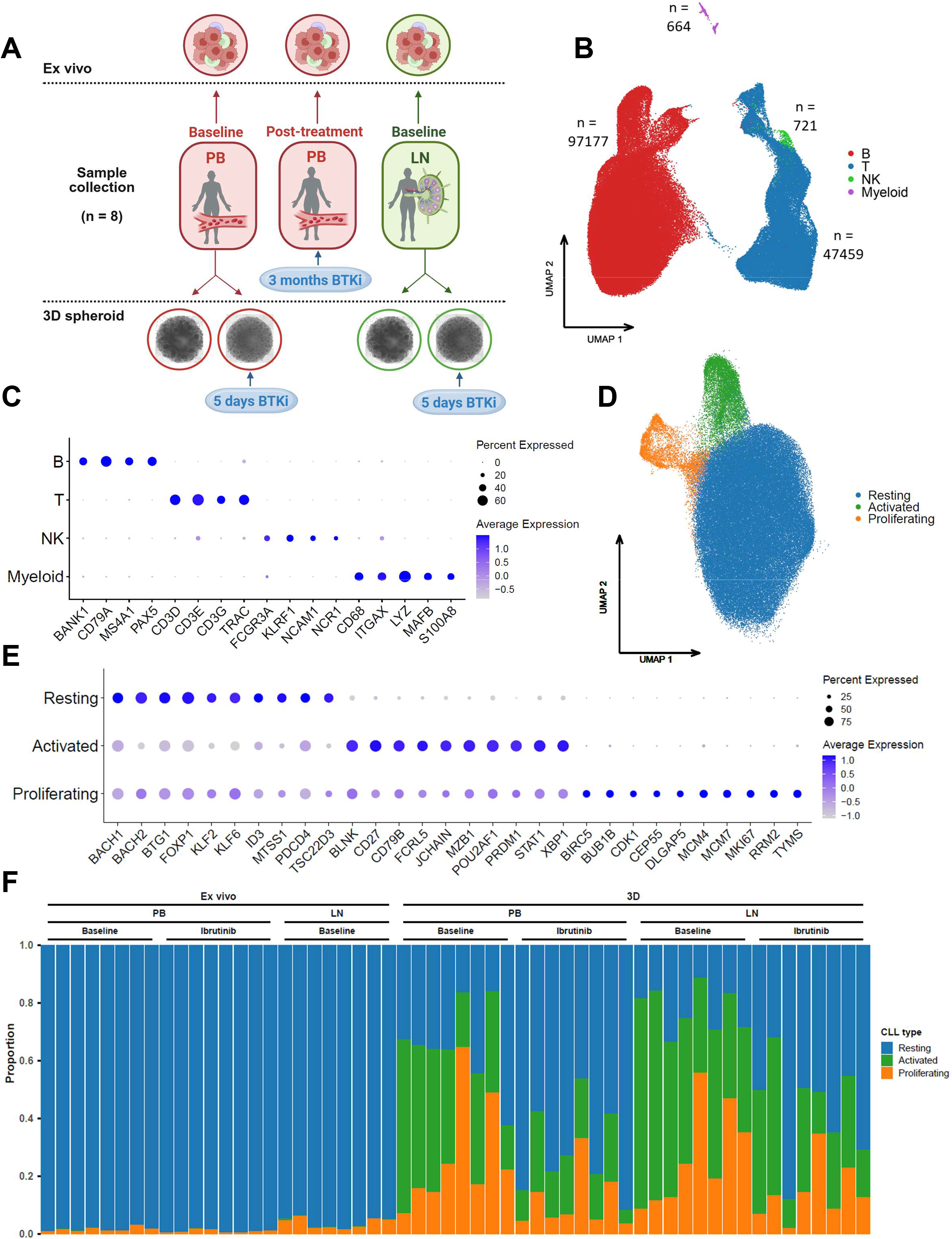
Single-cell sequencing reveals functional plasticity of CLL phenotypes upon ibrutinib treatment. A. Schematic overview of the study design. ScRNA-seq analysis was performed on paired PB and LN baseline samples of 8 CLL patients, including paired PB samples after 3 months of ibrutinib treatment. Baseline PB and LN samples were used to create 3D spheroids which were cultured for 5 days in vitro in the presence or absence of 100nM ibrutinib. Created in BioRender. B. Uniform manifold approximation and projection (UMAP) plot of 146,021 cells after integration of paired ex vivo PB, ex vivo LN, 3D PB and 3D LN samples before and after ibrutinib treatment from 8 CLL patients analyzed by scRNA-seq, identifying major immune cell populations annotated as B, T, NK and myeloid cells. C. Dot plot of the expression of marker genes in the 4 cell clusters. D. UMAP plot of 97,177 CLL cells identified in 1B analyzed by scRNA-seq, identifying 3 transcriptionally defined CLL states annotated as resting, activated and proliferating CLL cells. E. Dot plot of the expression of marker genes in the 3 CLL clusters. F. Stacked bar plot showing the proportions of resting, activated and proliferating CLL clusters out of total CLL cells across all experimental conditions. Each bar corresponds to a patient.

### Data pre-processing

Raw FASTQ files were processed using Cell Ranger v8.0.0 (10x Genomics) with the ‘cellranger multi’ pipeline, aligning reads to the GRCh38 2020-A reference genome with introns included to capture unspliced transcripts and to a custom multiplexing reference for the hashing antibodies. The pipeline performed alignment, barcode assignment, hashtag-based demultiplexing, and generation of gene expression matrices. Barcodes that could not reliably be assigned to a single cell were excluded from downstream analysis. Resulting filtered feature-barcode matrices were analyzed using *Scanpy* (v1.9.3) in Python (v3.8.1) and R (v4.3.0). Initial filtering removed cells with fewer than 200 total UMI counts using ‘sc.pp.filter_cells’. Remaining doublets were identified using *scDblFinder*^13^, with the expected doublet rate adjusted to account for prior removal of inter-sample doublets by hashtag-based demultiplexing, assuming approximately 1/7 of doublets remained after hashtag demultiplexing due to intra-sample doublets. Low-quality or outlier cells were identified using 4 QC metrics: log10-transformed total counts, log10-transformed number of detected genes, percentage of counts in the top 20 expressed genes, and mitochondrial content. Cells were flagged as outliers if they deviated by more than 5 median absolute deviations (MADs) from the median for the first 3 metrics, or had mitochondrial content exceeding 15%^14^. Cells classified as outliers by at least 2 of these metrics were excluded from downstream analysis. After cell filtering, genes expressed in fewer than 10 cells were removed. The resulting count data were normalized using median count depth normalization with log1p transformation, yielding a final dataset of 146,021 cells across 56 samples.

### Integration and annotation

The normalized data were further analyzed in R using *Seurat* (v5.1.0)^15^. Initial integration of the 112 samples (56 samples in duplicate) was performed using Seurat’s sketch-based reciprocal principal component analysis (RPCA) integration with 500 representative cells per sample. The sketch-based integrated reference was then used to integrate all cells into the reference space. Graph-based clustering was performed on the integrated data at multiple resolutions, and cells were manually annotated based on canonical marker expression to distinguish 4 major compartments: B, T, NK, and myeloid cells.

Next, the B cell and T/NK cell compartments were re-integrated separately. T/NK cells were integrated across the 7 conditions (that is, each unique combination of system, tissue, and treatment) using RPCA without sketching. B cells were integrated across the 56 biological replicates (that is, each unique combination of system, tissue, treatment, and patient) using sketch-based RPCA with 500 representative cells per replicate. Graph-based clustering was performed on the integrated B and T/NK cells data at multiple resolutions. The *clustree* R package^16^ was used to inform the choice of appropriate resolutions and identification of clusters. Genes differentially expressed between a cluster and all other clusters were determined at multiple resolutions using Seurat’s ‘FindAllMarkers’ function to support annotation. Manual annotation was refined for all cells using canonical markers and cluster-specific gene expression patterns.

### Compositional analysis

CLL B cells and T/NK cells were analyzed separately because they were physically sorted into distinct populations (60% CLL, 40% non-CLL) prior to sequencing. For statistical testing of compositional differences, we applied the ‘propeller’ framework from the *speckle* R package^17^. Cell type proportions were first logit-transformed to reduce heteroscedasticity, then empirical Bayes moderated t-tests were performed using the linear model-based approach of the *limma* R package (function ‘lmFit’). To account for within-patient correlation, we included the consensus correlation between samples from the same patient (function ‘duplicateCorrelation’). Resulting P-values were corrected for multiple testing using the Benjamini-Hochberg false discovery rate (FDR) method.

### Differential gene expression and gene set analysis

Genes differentially expressed between conditions for cells of the same cell type were determined using a pseudobulk-based analysis. Pseudobulk gene expression matrices were generated by aggregating raw counts using Seurat’s ‘AggregateExpression’ function, grouping cells by unique combination of system, tissue, treatment, patient, and cell type. Statistical analyses were performed separately per cell type using the *edgeR* and *limma* R packages. Lowly expressed genes were filtered out using ‘filterByExpr’ with a minimum count threshold of 10. Count data were transformed to log2-counts per million (logCPM), normalized by applying the trimmed mean of M-values (TMM) method (function ‘calcNormFactors’).

Precision weighting was applied using the voom method, and differential expression was assessed using empirical Bayes moderated t-tests incorporating these precision weights. A consensus intra-patient correlation was estimated for pseudobulk samples from the same patient (function ‘duplicateCorrelation’) and included in the linear model fit. Resulting P-values were corrected for multiple testing using the Benjamini-Hochberg FDR method.

Gene set enrichment analysis was performed using ‘camera’ from the *limma* R package, testing for enrichment of Hallmark and C2 gene sets from the Molecular Signatures Database (MSigDB; v2024.1) and curated gene sets retrieved from the literature. Gene symbols were harmonized using the *HGNChelper* R package to preserve already matched symbols while updating outdated ones. Gene set indices were constructed using ‘ids2indices’. P-values were determined using camera’s rank-based competitive test to assess directional enrichment (‘up’ or ‘down’) and corrected for multiple testing using the Benjamini-Hochberg FDR method. Results for selected gene sets and comparisons were visualized using heatmaps and enrichment networks. For the latter, the *aPEAR* R package was used to group similar gene sets together and to highlight the most important themes in the gene set enrichment results. Default settings for the calculation of the similarity matrix between gene sets (‘jaccard’), clustering (‘markov’), and naming of the clusters (‘pagerank’) were used.

T/NK cells were scored (*Seurat*; function ‘AddModuleScore’) for a curated list of cytotoxicity genes and the MSigDB HALLMARK_TNFA_SIGNALING_VIA_NFKB gene set. Module scores for each cell type were compared between lymph node and peripheral blood cells using one-sided Welch’s t-tests (LN < PB). P-values were corrected for multiple testing across cell types using the Benjamini-Hochberg FDR method. Results were visualized using violin plots with statistical significance indicated by asterisks.

### Cell-cell communication analysis

The *CellChat* R package was used for systematic analysis of cell-cell communication patterns using CellChatDB v2 as ligand-receptor interaction database. For the condition-specific analysis we followed the steps outlined in the package vignette ‘Inference and analysis of cell-cell communication using CellChat’ (https://htmlpreview.github.io/?https://github.com/jinworks/CellChat/blob/master/tutorial/CellChat-vignette.html). In summary, condition-specific CellChat objects were created from the integrated dataset including both B (CLL) and T/NK cells to enable cell-cell communication analysis between all cell types. Inferred communication probabilities based on ligand-receptor co-expression and associated p-values were calculated for all ligand-receptor interactions between all pairs of cell types using the function ‘computeCommunProb’ using the 5% truncated mean to calculate average expression. The function ‘netAnalysis_signalingRole_scatter’ was used to visualize the outgoing and incoming interaction strengths for each cell type per condition to highlight dominant senders (sources) and receivers (targets). Significant ligand-receptor interactions from T cell types to CLL cell types were extracted and visualized as bubble plots using the function ‘netVisual_bubble’.

For the analysis comparing cells before and after ibrutinib treatment, we followed the package vignette ‘Comparison analysis of multiple datasets using CellChat’ (https://htmlpreview.github.io/?https://github.com/jinworks/CellChat/blob/master/tutorial/Comparison_analysis_of_multiple_datasets.html). Condition-specific CellChat objects before and after ibrutinib treatment were merged. The information flow for each signaling pathway, that is the sum of the communication probabilities of the associated ligand-receptors across all pairs of cell groups, was calculated before and after treatment with T cell types as sources and CLL cell types as targets using the function ‘rankNet’ (mode=”comparison”). Signaling pathways were ranked based on the relative difference in the information flow between the two conditions. A paired Wilcoxon signed-rank test was performed to determine whether the difference in signaling information flow was statistically significant. Selected ligand-receptor pairs from altered signaling pathways were visualized as chord diagrams using the function ‘netVisual_chord_gene’.

## Results

### Single-cell sequencing reveals functional plasticity of CLL phenotypes upon ibrutinib treatment

To evaluate to what degree the 3D CLL spheroid model recapitulates the transcriptional states present in patients, scRNA-seq was performed on ex vivo CLL samples derived from matched PB and LN patient material, as well as in vitro spheroid (3D) cultures generated from these samples in the presence and absence of ibrutinib (Figure 1A). 8 treatment-naïve CLL patients were included of which 5 IGHV unmutated and 3 IGHV mutated (Supplemental table 1). Moreover, PB samples after 3 months ibrutinib treatment were collected to investigate to what extent the 3D ibrutinib response reflects the response in patients. Cells were sorted in a ratio of 60% CLL cells versus 40% non-CLL cells before proceeding with scRNA-seq.

After quality control, a total of 146,021 high-quality cells were obtained across 56 samples. Following data integration (Supplemental figure 1), major immune cell populations were identified by manual cell annotation, including CLL B cells, T cells, NK cells and myeloid cells (Figure 1B-C, Supplemental Data 1, Supplemental Figure 2). Next, we selected only CLL cells for analysis and identified 3 transcriptional CLL states by manual annotation: resting CLL, activated CLL and proliferating CLL (Figure 1D, Supplemental Figure 3-5, Supplemental Data 2). Resting CLL cells were annotated as such based on the higher expression of transcriptional repressors and genes involved in the maintenance of B cell quiescence (Figure 1E). Activated CLL cells were characterized based on the expression of genes linked to B cell activation and signaling, and proliferating CLL cells were identified based on the additional expression of proliferation genes.

Compositional analysis showed that the large majority of CLL cells in ex vivo samples belonged to the resting cluster independent of tissue origin or treatment, though within the CLL compartment, LN samples contained significantly more activated and proliferating CLL cells compared to PB (Figure 1F, Supplemental Figure 6, FDR<0.05). No significant differences regarding CLL composition were observed between IGHV unmutated and IGHV mutated patients (Supplemental Figure 7). The 3D model showed enrichment of activated and proliferating CLL fractions, allowing for more in-depth investigation of this clinically relevant niche, specifically by using these fractions as a proxy for the in vivo LN setting. PB and LN spheroids showed a similar composition of CLL fractions. In vitro ibrutinib treatment resulted in a decrease of activated and proliferating CLL cells and a concurrent increase of resting CLL cells in both PB and LN spheroids, suggesting a suppressive effect of ibrutinib treatment.

### Activation-dependent convergence of transcriptional tissue profiles highlights the importance of upstream signals for functional CLL state

To accurately compare ex vivo tissue-specific differences with those observed in 3D spheroids, we focused first on resting cells, which constitute the predominant population in ex vivo samples, as activated and proliferating CLL populations were too small in the ex vivo samples for direct comparisons with 3D samples. We first characterized tissue-specific differences within ex vivo samples. Direct comparison of gene expression showed that 300 genes were differentially expressed (FDR<0.05) between ex vivo PB and LN compartments, of which 162 were upregulated in LN-derived CLL cells (Figure 2A, Supplemental Data 3). Corresponding genes were mainly associated with proliferation and metabolism. Consistently, gene set enrichment analysis (GSEA) showed enrichment of multiple gene sets related to oxidative phosphorylation, proliferation and signal transduction in LN-derived CLL cells (Figure 2B, Supplemental Data 4).

**Figure 2.**
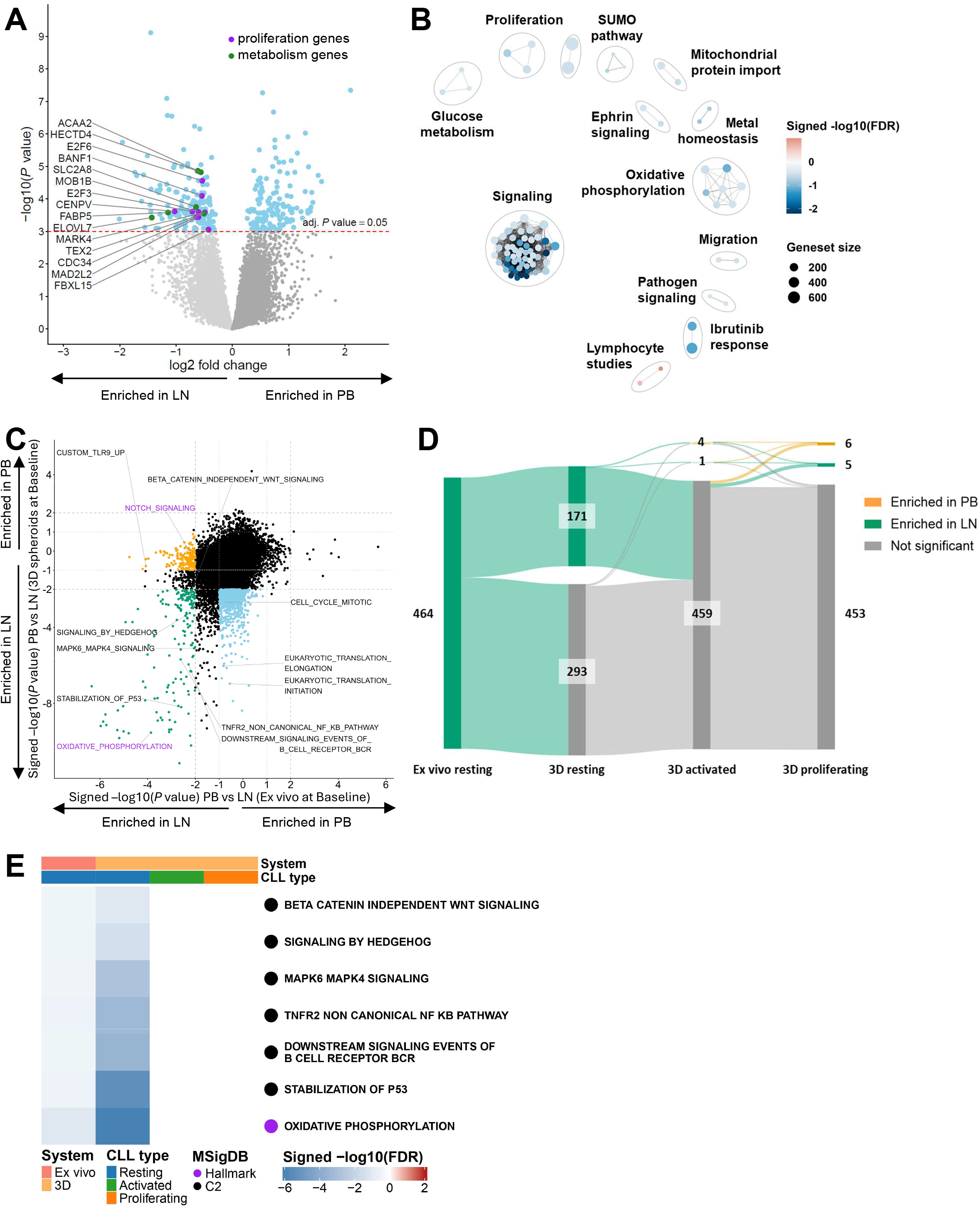
Activation-dependent convergence of transcriptional tissue profiles highlights the importance of upstream signals for functional CLL state. A. Volcano plot of log2 fold-change (FC) in gene expression between LN-derived (negative log-FC) and PB-derived (positive log-FC) CLL cells within ex vivo samples among resting CLL cells. Differentially expressed genes (FDR<0.05; n = 300) are highlighted in blue. Annotated genes indicate metabolism-related genes (highlighted in green) and proliferation-related genes (highlighted in purple). B. Enrichment map showing top 200 differentially enriched gene sets of LN-derived (blue) versus PB-derived (red) CLL cells within ex vivo samples, within the resting CLL cluster. Gene sets are grouped based on gene overlap, with edges connecting strongly overlapping gene sets. Annotation was added manually. C. Scatter plot of differentially enriched gene sets (P<0.01) of LN-derived (negative values) versus PB-derived (positive values) CLL cells within ex vivo samples (x-axis) and 3D samples (y-axis) among resting CLL cells. LN-enriched gene sets shared between ex vivo and 3D samples are highlighted in green. LN-associated gene sets significantly enriched only in ex vivo samples are highlighted in orange. LN-associated gene sets significantly enriched only in 3D samples are highlighted in blue. Annotations in purple indicate MSigDB hallmark gene sets and annotations in black indicate MSigDB C2 gene sets. D. Sankey plot of all gene sets enriched in LN-derived ex vivo resting CLL cells and whether they are enriched in either LN-derived or PB-derived CLL cells among the CLL clusters observed in 3D samples. ‘Not significant’ indicates that these genes sets are not differentially enriched between LN-derived and PB-derived CLL cells. E. Heat map showing differentially enriched gene sets of LN-derived (blue) versus PB-derived (red) CLL cells within all CLL clusters (ex vivo resting, 3D resting, 3D activated, 3D proliferating).

In order to investigate to what extent these tissue-specific features are conserved within 3D spheroids, we studied whether LN-associated gene sets in ex vivo samples were also enriched in LN spheroids compared to PB spheroids. Among resting CLL cells, gene sets enriched in LN-derived CLL cells across both ex vivo and 3D samples (highlighted in green) include BCR signaling, p53 signaling, TNFR-NF-κB signaling and oxidative phosphorylation (Figure 2C, Supplemental Data 5). LN-associated gene sets enriched uniquely in ex vivo samples (highlighted in orange) include TLR9 and Notch signaling, and LN-associated gene sets enriched uniquely in 3D samples (highlighted in blue) include translation and proliferation. Consistently, LN spheroids displayed higher in vitro CLL proliferation compared to PB spheroids (Supplemental Figure 8). In summary, direct comparison of resting CLL cells between ex vivo and 3D samples suggests conservation of transcriptional signatures associated with key signaling pathways previously described in CLL, such as BCR, p53 and NF-κB signaling^18,19^.

Since the 3D model is aimed at mimicking the clinically relevant CLL LN niche, we investigated whether CLL cells in the activated and proliferating clusters maintain the transcriptional profile of their tissue of origin. To this end, gene sets enriched in LN-derived cells in ex vivo samples were compared between LN and PB spheroids among the 3 different CLL populations. Whereas resting cells maintained some tissue identity at the transcriptional level, the majority of LN-enriched gene sets were not significantly enriched in either LN or PB spheroids among activated and proliferating CLL cells, suggesting that they converge to a similar phenotype (Figure 2D, Supplemental Figure 9). Consistently, several key pathways previously found to be enriched in LN-derived CLL cells in ex vivo samples showed similar activity between LN and PB spheroids among activated and proliferating CLL clusters (Figure 2E).

Altogether, these results support that upstream activation signals, rather than tissue of origin, are the primary determinants of functional CLL state, indicating that PB-derived CLL cells may be utilized to mimic a bona fide LN-activated CLL phenotype.

### 3D spheroids reveal divergent transcriptional responses to ibrutinib across CLL subpopulations

Next, the in vitro ibrutinib response in 3D spheroids was characterized and compared to the in vivo ibrutinib response in CLL patients. To this end, we compared ex vivo PB baseline and post-treatment samples of 8 CLL patients treated with ibrutinib monotherapy for 3 months. Corresponding PB and LN pre-treatment/baseline samples were used to create and culture 3D spheroids in the absence or presence of ibrutinib for 5 days. Ex vivo PB samples showed 491 downregulated (adjusted p<0.05, logFC<-1) and 253 upregulated (adjusted p<0.05, logFC>1) genes upon ibrutinib treatment in resting CLL cells (Figure 3A, Supplemental Data 6). Among the 3 CLL clusters in spheroids, resting CLL cells showed the highest number of differentially expressed genes upon ibrutinib treatment, whereas proliferating CLL cells showed the lowest number. Focusing on gene sets, PB and LN spheroids showed a strong correlation in downregulated (green) and upregulated (purple) gene sets upon ibrutinib treatment, indicating similar ibrutinib responses in PB and LN spheroids (Figure 3B, Supplemental Data 7). Notably, activated CLL cells exhibited the fewest downregulated gene sets, indicating that this population underwent the least pathway-level inhibition following ibrutinib treatment. In resting CLL cells, many pathways were significantly inhibited by ibrutinib, including BCR, TLR9, MYC, translation and proliferation (Figure 3C). Proliferating CLL cells exhibited a similar ibrutinib response. In contrast, the activated CLL fraction displayed a fundamentally distinct response to ibrutinib with no significant inhibition of any of the aforementioned pathways. Instead, activated CLL cells showed upregulation of TLR9, MALT1 and MYC pathways, as well as some translation and proliferation-associated gene sets upon ibrutinib treatment in specifically PB-derived spheroids. Notably, the IFNα pathway was upregulated upon ibrutinib treatment across all CLL subpopulations, which can promote CLL survival and is consistent with reports in ibrutinib-treated patients^20^.

**Figure 3.**
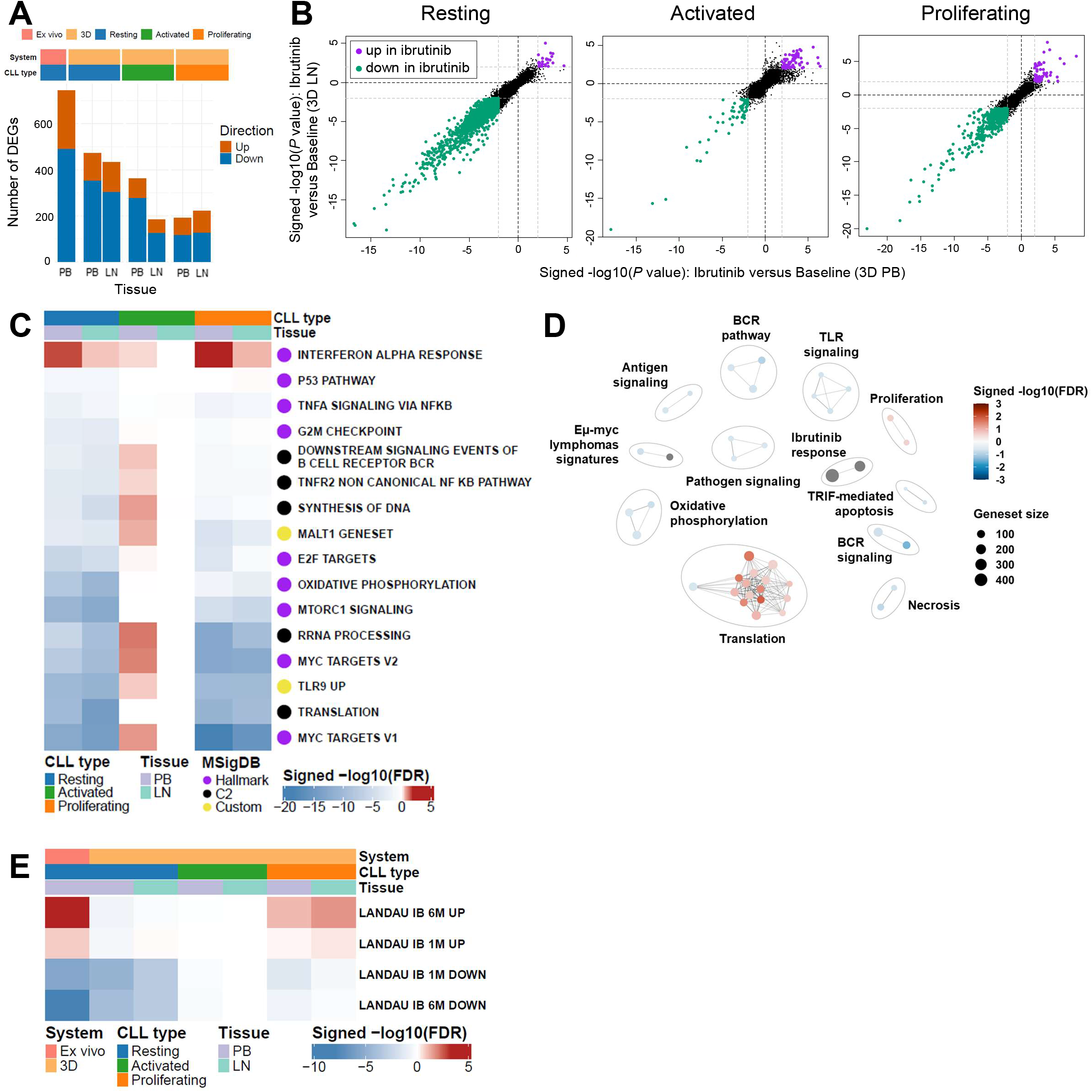
3D spheroids reveal divergent transcriptional responses to ibrutinib across CLL subpopulations. A. Stacked bar plot showing the number of significantly upregulated and downregulated genes (FDR<0.05; log2FC>1) upon ibrutinib treatment across all CLL clusters (ex vivo PB, 3D resting, 3D activated, 3D proliferating). B. Scatter plots showing upregulated and downregulated gene sets (P<0.01) upon ibrutinib treatment in PB spheroids (x-axis) and LN spheroids (y-axis) within all CLL clusters (resting, activated, proliferating). Shared upregulated gene sets are highlighted in purple. Shared downregulated gene sets are highlighted in green. C. Heat map showing upregulation (red) and downregulation (blue) of gene sets upon ibrutinib treatment within 3D PB and 3D LN samples across all CLL clusters (resting, activated, proliferating). D. Enrichment map showing upregulated (red) and downregulated (blue) gene sets (FDR<0.25) upon ibrutinib treatment in CLL cells within ex vivo PB samples, within the resting CLL cluster. Gene sets are grouped based on gene overlap, with edges connecting strongly overlapping gene sets. Annotation was added manually. E. Heat map showing upregulation (red) and downregulation (blue) of custom gene sets based on ibrutinib signatures from the dataset by Landau et al.^21^ upon ibrutinib treatment across all CLL clusters (ex vivo resting, 3D resting, 3D activated, 3D proliferating).

In order to find parallels between the in vitro ibrutinib response in spheroids and the ibrutinib response in patients, GSEA was performed on ex vivo samples comparing PB samples at baseline versus 3 months ibrutinib treatment. This indicated a similar downregulation of BCR and TLR signaling as well as oxidative phosphorylation (Figure 3D, Supplemental Data 8). Similar to 3D spheroids, upregulation of translation pathways was observed.

Finally, the ibrutinib response in CLL spheroids was compared to the ibrutinib response in patients, to which end custom gene signatures were created based on a previous bulk RNA sequencing study of PB-derived CLL cells from 14 ibrutinib-treated patients at 1 and 6 months of treatment^21^ (Supplemental Data 9). GSEA indicated that the response in our 3 month ibrutinib-treated patient cohort was consistent with both 1 month and 6 month time points reported previously^21^, showing similarity in both upregulated and downregulated genes (Figure 3E). Within spheroids, resting CLL cells showed similar downregulation of genes as observed in ibrutinib-treated patients. Proliferating CLL cells displayed similar downregulation of patient-downregulated genes alongside recapitulating upregulation of genes as observed in patients. Activated CLL cells showed the lowest similarity to the reported ibrutinib response^21^, consistent with the observation that this CLL subpopulation exhibited minimal inhibition by ibrutinib at the pathway level. Collectively, these data suggest lower sensitivity of activated and proliferating CLL cells to ibrutinib.

### 3D spheroids recapitulate tissue-associated T cell transcriptional features

We next focused on the T cell compartment in our samples, isolating all T cell clusters for subsequent reintegration. Unsupervised clustering resulted in 11 T cell clusters distinguished by manual annotation: 6 CD4 T cell clusters and 5 CD8 T cell clusters, as well as an NK cell cluster (Figure 4A, Supplemental Figure 10-11, Supplemental Data 10). The CD4 T cell clusters comprised of naïve, memory and effector differentiation states, as well as follicular and regulatory subsets. CD8 T cell clusters comprised of naïve, memory and EMRA differentiation states, as well as a separate proliferative CD8 T cell cluster. An additional T cell cluster with high expression of interferon-stimulated genes (ISGs) could further be distinguished in CD4 and CD8 T cells, annotated as CD4 ISG^+^ and CD8 ISG^+^. Finally, a cluster of NK cells was identified based on the unique expression of NK genes (Figure 4B). Similar to ex vivo samples, PB spheroids were enriched in CD8 TEMRA cells and LN spheroids were enriched in CD4 follicular and regulatory T cells (Figure 4C, Supplemental Figure 12, FDR<0.001). 3D spheroids were characterized by a prominent population of CD8 proliferating T cells independent of tissue origin, whereas the CD8 ISG^+^ population observed ex vivo was nearly absent in 3D spheroids. Apart from these populations, the in vitro 3D culture system had no substantial impact on T cell differentiation states. Ibrutinib treatment had no significant effect on T cell compositions in vitro, but did result in a reduced proportion of CD4 regulatory T cells in PB ex vivo.

**Figure 4.**
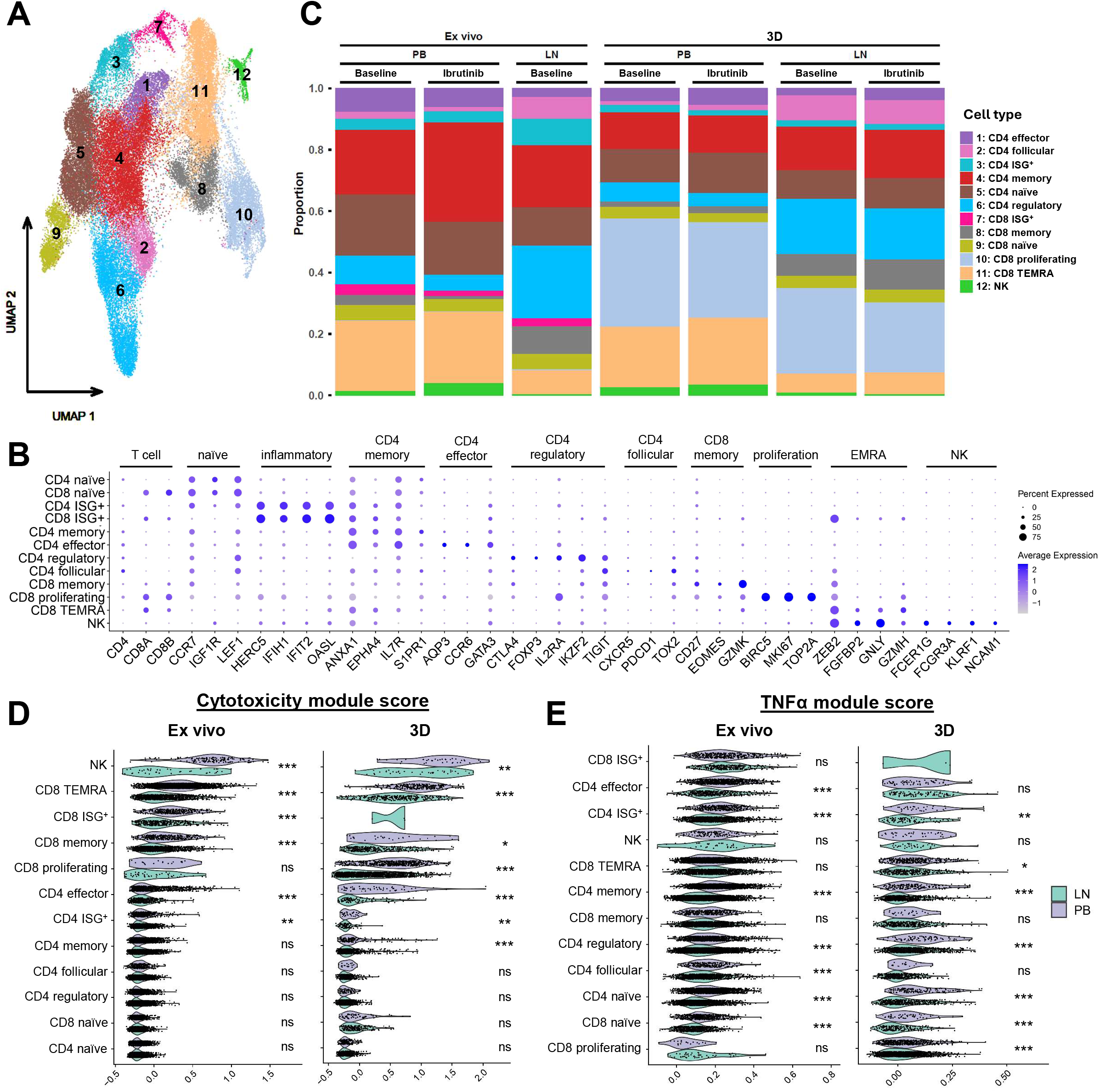
3D spheroids recapitulate tissue-associated T cell transcriptional features. A. UMAP plot of 48,180 T and NK cells after integration of paired ex vivo PB, ex vivo LN, 3D PB and 3D LN samples before and after ibrutinib treatment from 8 CLL patients analyzed by scRNA-seq, identifying 12 clusters divided into 6 CD4+ T cell clusters, 5 CD8+ T cell clusters and 1 NK cell cluster. B. Dot plot of the expression of marker genes in the 12 cell clusters. C. Stacked bar plot showing the proportions of T and NK cell clusters out of total T and NK cells across all experimental conditions. Each bar corresponds to the average proportions over 8 patients. D. Violin plots showing module scores of a cytotoxicity-associated gene signature^49^ in LN-derived versus PB-derived T cells within ex vivo and 3D conditions (baseline). E. Violin plots showing module scores of a TNFα gene signature (MSigDB hallmark) in LN-derived versus PB-derived T cells within ex vivo and 3D conditions (baseline).

For both ex vivo and 3D samples, gene signature scoring showed higher T cell cytotoxicity scores of PB-derived T cells compared to T cells from LNs within CD8 TEMRA and memory T cells, CD4 effector and ISG^+^ T cells as well as NK cells (Figure 4D). Similarly, CD4 ISG^+^, memory and regulatory T cells as well as naïve CD4 and CD8 T cells derived from PB showed higher levels of TNFα pathway activity compared to T cells from LNs in both ex vivo and 3D samples (Figure 4E). These data suggest that PB-derived T cells exhibit a more cytotoxic and pro-inflammatory phenotype, whereas LN-derived T cells display a more regulatory profile. In conclusion, these data revealed distinct tissue-specific functional T cell profiles that are largely preserved in 3D CLL spheroids.

### Cell-cell communication analysis suggests T cell-mediated support of the CLL LN niche and distinct **immunomodulatory** effects of ibrutinib

To better understand the role of T cells in supporting the CLL LN niche and the impact of ibrutinib on their intercellular communication, we next investigated CLL-T cell communication. To this end, cell-cell communication networks were characterized by predicting ligand-receptor pairs based on a curated literature-supported database by applying CellChat v2^22^. First, we used ex vivo LN samples as a benchmark and to assess consistency with prior studies^23^. In ex vivo LN samples, CD4 T cell populations were predicted to have the lowest degree of cell-cell communication, CD8 T cell populations to have the highest incoming interaction strength and CLL populations to have the highest outgoing one (Figure 5A), in line with recent analyses^23^. Furthermore, among CLL subclusters, proliferating CLL cells were predicted to have the highest outgoing and activated CLL cells the highest incoming interaction strength. The vast majority of CLL-to-T cell interactions previously predicted to be present in CLL LN samples^23^ were also inferred in the present analysis (Supplemental Figure 13). With regards to T-to-CLL cell interactions, the majority of predicted ligand-receptor pairs in ex vivo LN samples showed concordant predictions in 3D spheroids (Figure 5B, Supplemental figure 14). Corresponding ligand-receptor pairs in this group included interactions associated with signaling pathways known to contribute to CLL support including TNFR^24^, TGF-β^23^, MIF^25^ and CD40^26^.

**Figure 5.**
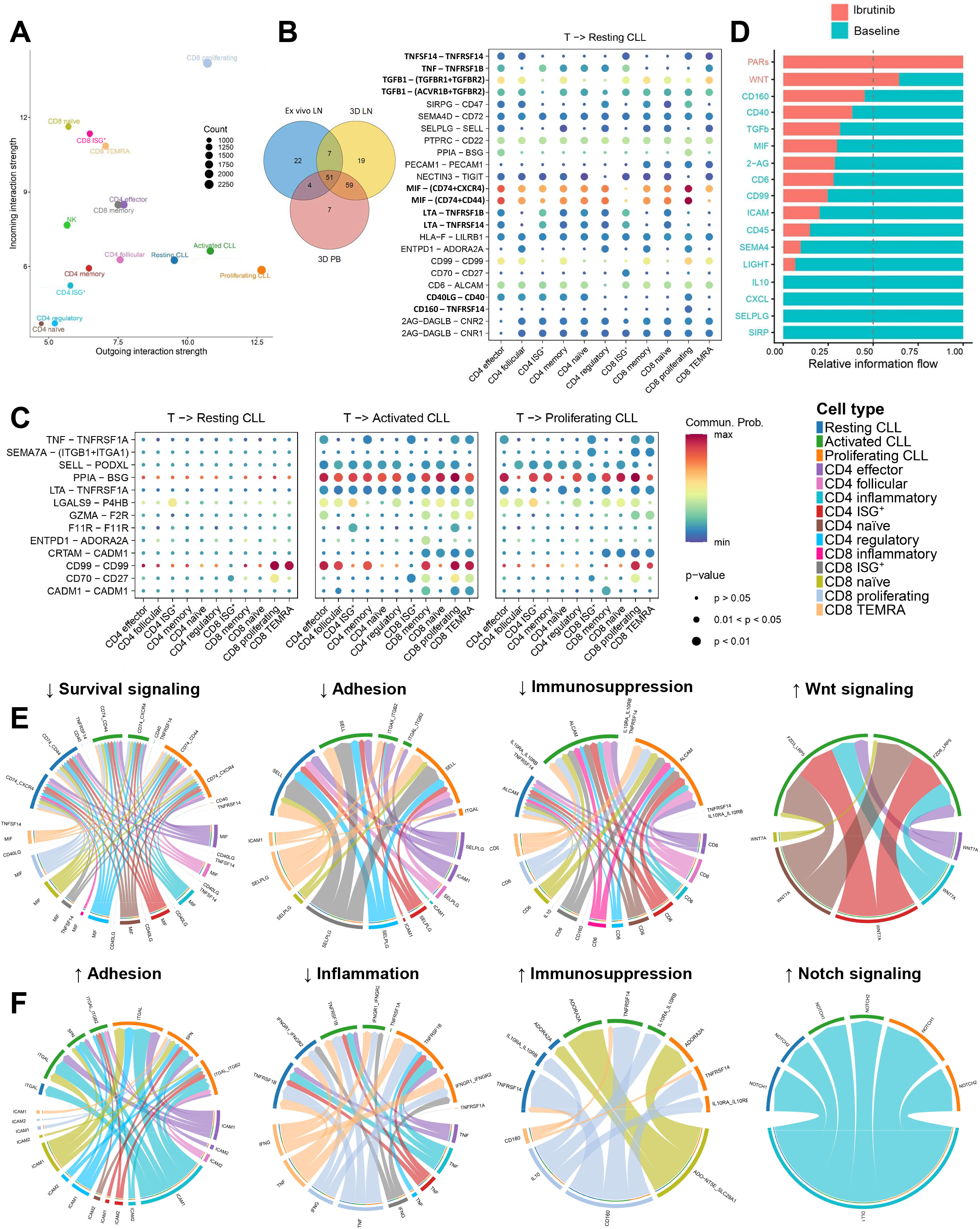
Cell-cell communication analysis suggests T cell-mediated support of the CLL LN niche and distinct immunomodulatory effects of ibrutinib. A. Scatter plot showing outgoing interaction strength (x-axis) and incoming interaction strength (y-axis) of each annotated cell cluster in ex vivo LN baseline samples. B. Venn diagram showing overlap of predicted T-to-CLL ligand-receptor pairs in steady state analysis of ex vivo LN, 3D LN and 3D PB conditions at baseline. Corresponding bubble plot shows selection of these shared T-to-CLL ligand-receptor pairs within ex vivo LN baseline samples among resting CLL cells. The dot color and size represent the calculated inferred communication probability and p-values. C. Bubble plots showing selection of predicted T-to-CLL ligand-receptor pairs within 3D LN baseline samples that are shared between 3D LN and 3D PB conditions that engage activated and proliferating CLL cells while remaining inactive in resting CLL cells. The dot color represents the calculated inferred communication probability and is relative and can thus not be compared between bubble plots. The dot size represents the calculated inferred p-values. D. Stacked bar plot showing selection of rankNet analysis comparing baseline versus ibrutinib in ex vivo PB samples. Significant signaling pathways are ranked based on differences in overall information flow within the inferred networks between the two conditions. E. Chord plots of up-/downregulated T-to-CLL interactions upon ibrutinib treatment in ex vivo PB samples. The outer bars indicate the involved cell populations. The inner bar size is proportional to the signal strength received by the target. F. Chord plots of up-/downregulated T-to-CLL interactions upon ibrutinib treatment in 3D PB spheroids. The outer bars indicate the involved cell populations. The inner bar size is proportional to the signal strength received by the target.

To investigate which T-to-CLL cell interactions are associated with activated and proliferating CLL states, further analysis focused on the 110 interactions shared between LN and PB spheroids. We selected interactions in LN spheroids that specifically engaged activated and proliferating CLL populations while remaining inactive in resting CLL cells (Figure 5C, Supplemental Figure 15). Several inferred interactions were related to signaling via TNFR1^24^ and PPIA-BSG^27^, as well as cell-cell binding via integrins and cell adhesion molecules, in line with the central role of inflammatory pathways and microenvironmental support in the activation and proliferation of CLL cells^28^.

Next, we performed differential CellChat analysis to explore how 3 months ibrutinib treatment may affect T-to-CLL cell communication. By quantifying communication probabilities of ligand-receptor pairs within signaling networks across samples, this approach aims to identify which pathways may be affected upon ibrutinib treatment. Based on ex vivo PB samples, computational predictions indicated that ibrutinib treatment downregulates several pathways involved in CLL support (CD40, MIF, CD45, LIGHT) and adhesion (CD99, ICAM, SELPLG) (Figure 5D, Supplemental Figure 16). Additionally, downregulation of several immunosuppressive pathways (TGFb, CD6, IL10, SIRP) was predicted, pointing to immunomodulatory effects of ibrutinib. Notably, increased Wnt pathway activity was predicted, suggesting a potential mechanism by which CLL cells may be able to persist during BTK inhibition.

Zooming in on specific T-to-CLL ligand-receptor pairs, ibrutinib was predicted to disrupt multiple inferred T-CLL cell interactions, among which CD40, MIF-CD74 and TNFRSF14, reflecting a downregulation of multiple signaling networks that sustain CLL support (Figure 5E, Supplemental Figure 17). Several interactions were involved in adhesion, such as SELPLG and ICAM1, the latter exclusively in activated and proliferating CLL populations. Ibrutinib treatment was predicted to downregulate several immunosuppressive mediators such as CD6, IL10 and CD160, suggesting reshaping of the PB microenvironment upon treatment. Notably, increased Wnt signaling was predicted via T cell-derived WNT7A engaging FZD3/6 on specifically activated CLL cells, suggesting a possible association with this phenotype.

Finally, we performed the same analysis in ibrutinib-treated PB spheroids, to investigate how ibrutinib may affect T-CLL cell communication within the LN niche. In contrast to ex vivo PB samples, ibrutinib treatment of 3D spheroids was inferred to result in increased ICAM1-mediated adhesion, suggesting increased predicted adhesion-related signaling (Figure 5F). Consistently, no significant predicted inhibition of survival-associated pathways was observed in 3D spheroids. Moreover, in vitro ibrutinib treatment was inferred to result in upregulation of immunosuppressive mediators such as IL10 and downregulation of pro-inflammatory mediators such as TNF, IFN-γ and LT-α. This suggests an immunomodulatory effect opposite to that predicted in ex vivo PB samples, toward an immunosuppressive state. Finally, increased Notch signaling was predicted via T-cell derived DLL1 engaging NOTCH1/2 on CLL cells, representing a potential survival mechanism under ibrutinib treatment. Collectively, computational cell-cell communication analyses support T cell-mediated interactions contributing to the LN niche and predict niche-dependent differences in T-CLL signaling interactions.

## Discussion

The LN is the critical site of CLL proliferation and development of resistance to therapeutic agents^29,30^. Previous scRNA-seq studies distinguished 3 major CLL cell states, quiescent, activated and proliferating, yet showed that the activated CLL cells accounted for only 2.2-4.3% in LN samples and proliferating CLL cells comprised 0.4-1.0% of cells^18^. Here, we utilized our established 3D CLL spheroid model^10^ to increase the proportion of activated and proliferating CLL cells, enabling analysis of these otherwise rare populations in an in vitro system capturing aspects of the LN microenvironment. This 3D system is relatively simple and does not contain e.g. stromal cells and a minimal myeloid component. Presence of these cells in patient LNs is variable and to include them is logistically and experimentally complex, therefore we have chosen a minimal and reproducible setup which reflects important aspects of CLL pathobiology, as discussed below.

To benchmark physiological relevance of the 3D model in mimicking a LN phenotype, tissue-specific differences were identified in ex vivo LN and PB samples, in which the majority of CLL cells displayed a resting phenotype. Whereas resting CLL cells within spheroids indeed partly maintained these transcriptional differences, CLL cells within the activated and proliferating subpopulations showed similar activity of corresponding pathways independent of tissue origin, indicating that PB-derived CLL cells can adopt transcriptional features associated with LN-activated CLL states under these culture conditions.

With regards to divergent CLL phenotypes, we observed a shift of activated and proliferating CLL cells to a resting state upon ibrutinib, consistent with the reported acquisition of a quiescence-like phenotype in CLL cells under ibrutinib treatment over time^31^. Deeper investigation of the ibrutinib response in CLL spheroids revealed divergent responses across CLL subpopulations. Resting and proliferating CLL cells were similarly affected by ibrutinib characterized by the inhibition of many key pathways as well as significantly increased IFNα pathway activity. This suggests that when BTK is blocked, CLL cells show transcriptional signatures consistent with activation of compensatory signaling pathways. A potential explanation is TLR-mediated induction of an autocrine IFNα feedback loop shown to work independent of BTK^32,33^, which fits with the stronger IFNα induction in PB spheroids that displayed higher TLR9 signaling capacity. This mechanism is consistent with persistent IFN signaling previously observed in ibrutinib-treated patients^20^ and synergy of ibrutinib with TLR9 antagonists observed in multiple studies^34,35^. Combined with the fact that multiple studies showed the relevance of TLR9 signaling in the CLL LN niche^6,19,30,34^, these findings underscore TLR9 signaling as a candidate pathway that may contribute to differential responses to ibrutinib and warrants further investigation.

Whereas resting and proliferating CLL cells were effectively targeted by ibrutinib, the activated CLL cell population showed predominant refractoriness to ibrutinib, concurrent with MYC-mediated activation of translation and proliferation pathways. This induction was observed specifically in PB spheroids together with increased TLR9 and MALT1 pathway activity. In agreement, MALT1 has been shown to be able to directly regulate MYC protein expression, thereby controlling MYC-driven gene expression networks^36,37^. Notably, MALT1 is downstream of BTK in the BCR signaling pathway and its activity can bypass BTK inhibition, representing a possible route to ibrutinib resistance^38,39^. The concurrent increase of TLR9 pathway activity, points toward a TLR9-MALT1-MYC signaling axis. A potential explanation for PB spheroids displaying a greater capacity to trigger a robust TLR9 response within 3D cultures is that LN-derived CLL cells have higher baseline TLR9 signaling^34^, leading to receptor desensitization. TLR9-mediated upregulation of MYC and mTOR pathways was previously described in CLL cells together with upregulation of BCR signaling components^40^. A previous scRNA-seq study on CLL PBMCs revealed a subpopulation of CLL cells with increased expression of MYC targets and mitotic cell genes, reflecting circulating LN-activated CLL cells^41^. This CLL subpopulation was highly expanded upon progression during ibrutinib, suggesting a pre-existing threat of relapse. Notably, following ibrutinib treatment in our patient samples, PB-derived CLL cells showed similar upregulation of translation pathways but without upregulation of proliferation pathways, suggesting that PB CLL cells in vivo are similarly primed for proliferation, but lack additional upstream activation signals to actually induce it. A previous scRNA-seq study on LN-derived CLL has suggested that CLL cells start in the proliferating state, move to the activated state and end in the quiescent state based on RNA velocity analysis^18^. Our data based on the spheroid system suggest a more dynamic context in which activated cells retain the potential to induce proliferation when subjected to ibrutinib drug pressure. Even though the proliferative CLL fraction is considered as the main contributor to disease progression, these new data suggest that the activated CLL fraction may pose a bigger threat to relapse.

A key difference between LN and PB samples used to create CLL spheroids is the higher proportion of CD4 T cells in LN samples (Supplemental Figure 18). Higher activity of proliferation pathways was observed in LN spheroids, supporting the role of T cells in promoting CLL proliferation^10^. This increased LN CLL proliferation was maintained upon ibrutinib treatment (data not shown), suggesting that the cellular microenvironment may protect the CLL LN niche during ibrutinib treatment. Consistent with previous studies^23,42^, we observed a more cytotoxic phenotype in PB-derived T cells and a more immunoregulatory phenotype in LN-derived T cells. Despite these tissue-specific T cell phenotypes, PB and LN spheroids largely resembled each other in terms of ibrutinib response and T-to-CLL cell communication. With respect to this, T-CLL cell communication analysis predicted tissue-specific immunomodulation upon ibrutinib treatment, in which ex vivo PB samples show a shift to a less immunosuppressive state whereas LN niche-mimicking 3D spheroids shifted to a more immunosuppressive state. This was corroborated by a predicted interruption of interactions mediating cell-cell adhesion in PB samples, whereas adhesion in 3D spheroids increased upon ibrutinib. Clinically, this may imply that niche-derived cues may cooperatively sustain CLL and mediate resistance to therapy. In 3D spheroids, ibrutinib treatment resulted in the prediction of increased T-to-CLL interactions involved in Notch signaling. This pathway has been described to mediate CLL progression in mouse models^43^ and *NOTCH1* mutations have frequently been reported in CLL and Richter’s transformation to drive disease progression^44^. In vivo studies based on PB samples of ibrutinib-treated CLL patients reported diminished immunosuppressive properties (i.e. decreased regulatory T cell numbers and reduced expression of inhibitory receptors) and increased T cell effector functions after treatment^45–48^, consistent with our cell-cell communication analyses. While our cell-cell communication analysis provides predictive insights into potential interactions within the CLL LN niche, they remain unvalidated hypotheses that require experimental confirmation.

In summary, this work demonstrates that 3D CLL spheroids capture valuable transcriptional features of CLL heterogeneity and drug responses observed in vivo. In addition to coexistence and differential responses of resting, activated, and proliferating CLL phenotypes, the single-cell data identify a persistent, ibrutinib-insensitive activated CLL fraction. This subset displays unique transcriptional adaptations which may be leveraged to guide future therapeutic interventions overcoming microenvironment-induced resistance.

### Limitations of the study

A number of limitations should be acknowledged. First, the 3D spheroid system relies on stimulation with IL-2, IL-15, IL-21, and CpG, which is likely to affect transcriptional programs and may confound interpretation of cell states, pathway activity, and drug responses relative to the native LN microenvironment. Second, cells were artificially reconstituted prior to sequencing at a fixed 60:40 ratio of CLL to non-CLL cells. While this does not bias within-compartment analyses, it precludes direct comparison of overall cellular composition across all cell types and requires interpretation of compositional and cell–cell communication analyses to be restricted within defined compartments.

Third, this study is based primarily on transcriptomic profiling of *ex vivo* patient-derived samples and paired *in vitro* 3D cultures. As a result, the proposed resistance mechanisms, predicted signaling pathway activities, and inferred cellular interactions should be interpreted with caution, positioning this work as descriptive and hypothesis-generating rather than providing definitive mechanistic conclusions. The relatively small cohort size of 8 patients limits broader generalizability. Finally, the 5-day in vitro ibrutinib exposure captures only short-term responses and cannot fully model the long-term evolutionary processes underlying clinical resistance.

### Author contributions

MVH designed the study, performed experiments, analyzed data, guided figure design and edited figures, and wrote the paper. DL designed the study, performed bioinformatic analyses, generated figures, and wrote the paper. AJ performed bioinformatic analyses and generated figures. JLR and MD performed experiments. JD managed patient data administration and coordinated sample selection. FSP guided data analysis and figure design, and corrected the paper. APK provided clinical expertise, patient samples and corrected the paper. PDM designed the study, guided data analysis and figure design, performed bioinformatic analyses, generated figures, and wrote the paper. EE conceptualized the study, guided data analysis and figure design, provided budget support (MVH, DL, JLR), and wrote the paper.

## Acknowledgements

Remco Loos (Center for Innovation and Translational Research Europe, Bristol Myers Squibb, Sevilla, Spain) and Jeroen Bakker (Acerta Pharma B.V., Oss, The Netherlands) were involved in the development of the CLOSIT project. We used patient samples from the HOVON 158 trial and we thank the corresponding clinicians, trial staff and patients involved.

