## Supplemental Table 1 for "Single-cell characterization of 3D CLL spheroid model that recapitulates lymph node microenvironment-driven phenotypes and in vivo drug responses to ibrutinib"

| sample ID | tissue | treatment | sample date | age | gender | IGHV | %MRD | WBC (G/L) | %lymphocytes | FISH | TP53 |
|---|---|---|---|---|---|---|---|---|---|---|---|
| H158-002 | PB | none | 4-1-2021 | 52 | M | mutated | 93,27 | 179,83 | 88,4 | del(13q14) | wild type |
| H158-002 | LN | none | 4-1-2021 | 52 | M | mutated | - | - | - | del(13q14) | wild type |
| H158-002 | PB | ibrutinib | 30-3-2021 | 52 | M | mutated | 56,56 | 22,13 | 76,3 | del(13q14) | wild type |
| H158-005 | PB | none | 20-1-2021 | 72 | M | unmutated | 88,39 | 105,75 | 91,1 | del(13q14) | wild type |
| H158-005 | LN | none | 20-1-2021 | 72 | M | unmutated | - | - | - | del(13q14) | wild type |
| H158-005 | PB | ibrutinib | 23-4-2021 | 72 | M | unmutated | 82,47 | 17,21 | 77,4 | del(13q14) | wild type |
| H158-036 | PB | none | 25-3-2021 | 75 | M | mutated | 89,74 | 99,78 | 83 | del(13q14) | wild type |
| H158-036 | LN | none | 25-3-2021 | 75 | M | mutated | - | - | - | del(13q14) | wild type |
| H158-036 | PB | ibrutinib | 17-6-2021 | 75 | M | mutated | 76,49 | 91,92 | - | del(13q14) | wild type |
| H158-038 | PB | none | 31-3-2021 | 73 | M | mutated | 94,88 | 331,56 | 91,1 | del(11q), del(13q14) | wild type |
| H158-038 | LN | none | 31-3-2021 | 73 | M | mutated | - | - | - | del(11q), del(13q14) | wild type |
| H158-038 | PB | ibrutinib | 22-6-2021 | 73 | M | mutated | 93,88 | 169,28 | 93,6 | del(11q), del(13q14) | wild type |
| H158-039 | PB | none | 31-3-2021 | 63 | F | unmutated | 82,37 | 22,61 | 89,7 | tri12 | wild type |
| H158-039 | LN | none | 31-3-2021 | 63 | F | unmutated | - | - | - | tri12 | wild type |
| H158-039 | PB | ibrutinib | 23-6-2021 | 63 | F | unmutated | 44,21 | 15,15 | 86,5 | tri12 | wild type |
| H158-046 | PB | none | 11-5-2021 | 72 | F | unmutated | 90,63 | 105,81 | 92,95 | del(13q14) | wild type |
| H158-046 | LN | none | 11-5-2021 | 72 | F | unmutated | - | - | - | del(13q14) | wild type |
| H158-046 | PB | ibrutinib | 3-8-2021 | 72 | F | unmutated | 54,4 | 40,98 | 85,8 | del(13q14) | wild type |
| H158-056 | PB | none | 28-5-2021 | 45 | F | unmutated | 81,19 | 137,49 | 82 | del(13q14) | wild type |
| H158-056 | LN | none | 28-5-2021 | 45 | F | unmutated | - | - | - | del(13q14) | wild type |
| H158-056 | PB | ibrutinib | 25-8-2021 | 45 | F | unmutated | 86,65 | 67,68 | 92,1 | del(13q14) | wild type |
| H158-076 | PB | none | 24-6-2021 | 56 | M | unmutated | 85,09 | 460,26 | 89,2 | del(13q14) | wild type |
| H158-076 | LN | none | 24-6-2021 | 56 | M | unmutated | - | - | - | del(13q14) | wild type |
| H158-076 | PB | ibrutinib | 21-9-2021 | 56 | M | unmutated | 89,38 | 306,78 | 96,5 | del(13q14) | wild type |
