## Supplementary figures and images for "Single-cell characterization of 3D CLL spheroid model that recapitulates lymph node microenvironment-driven phenotypes and in vivo drug responses to ibrutinib"

### Supplemental Figure 1

#### Supplemental Figure 1

A

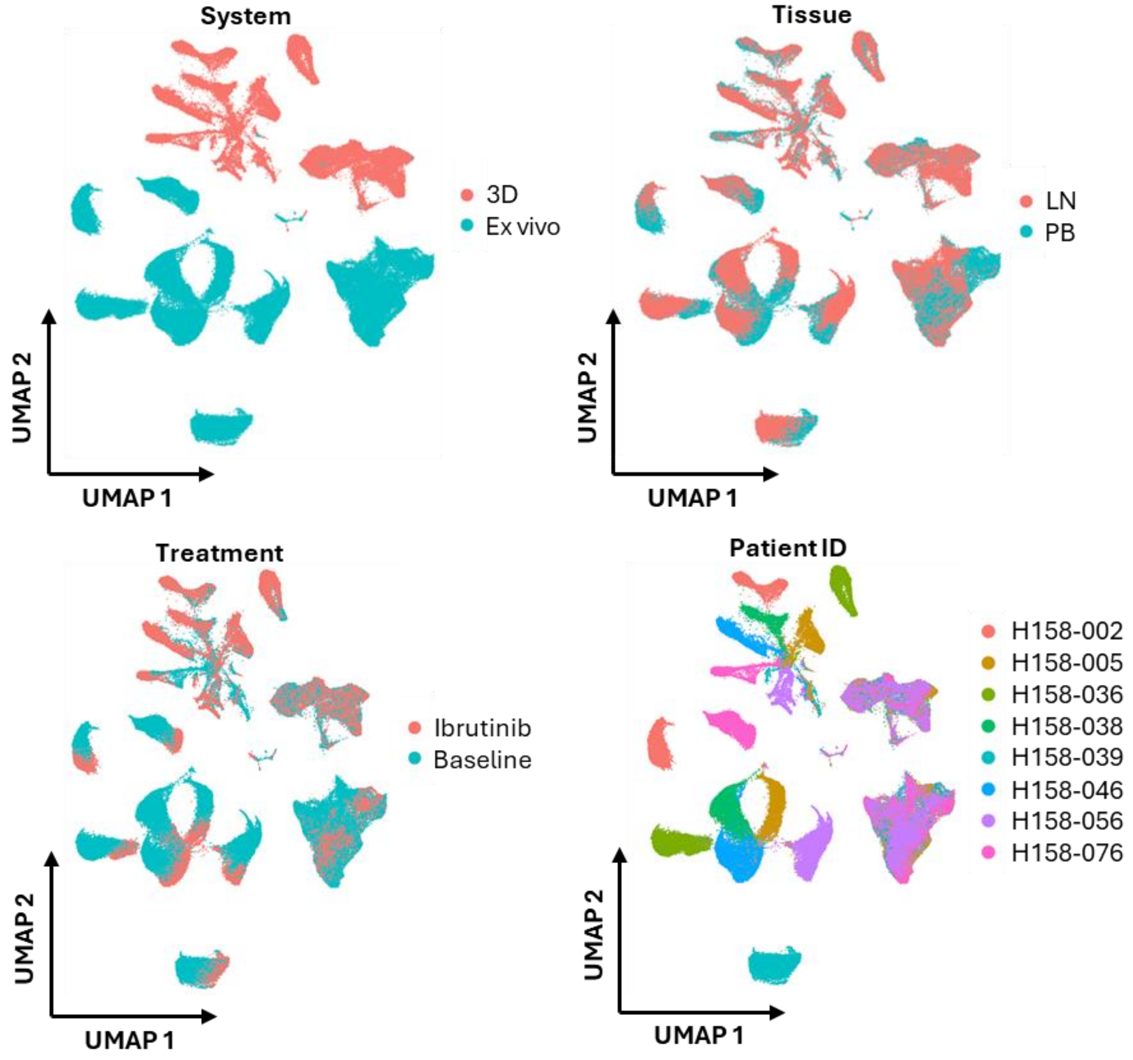


B

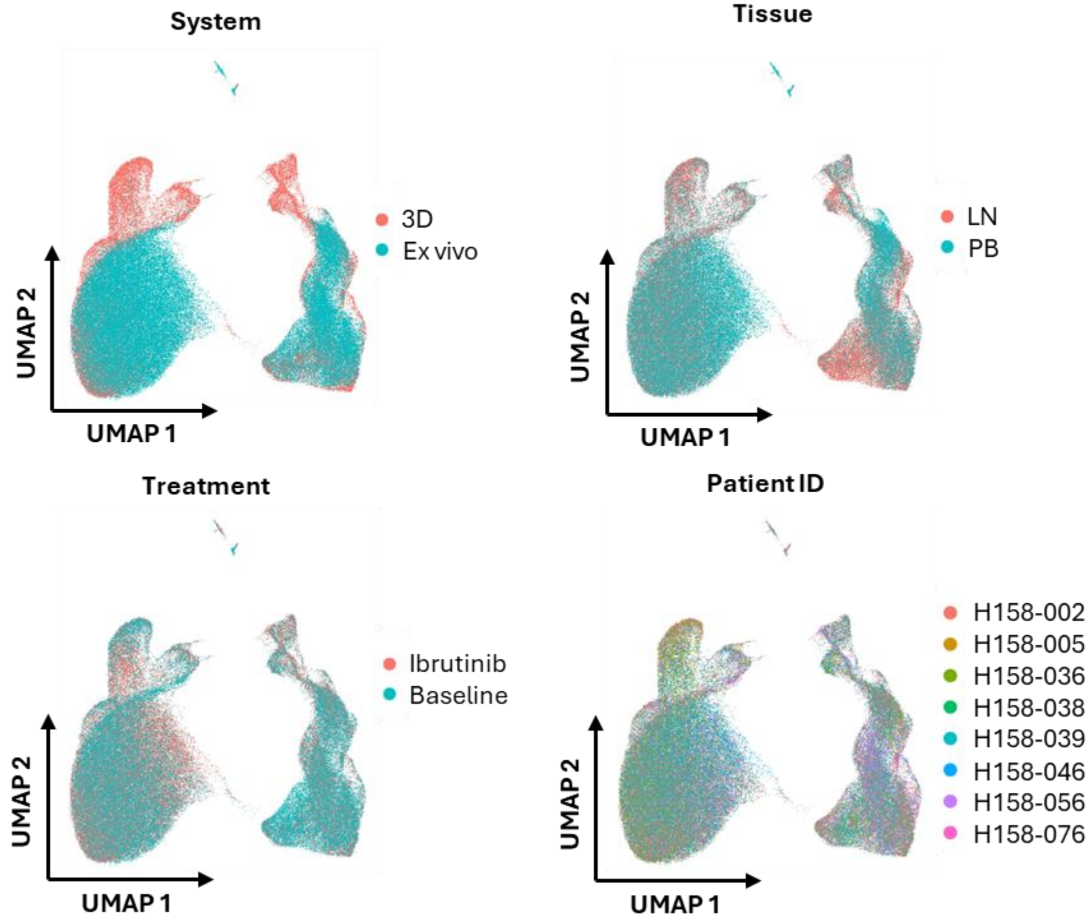

### Supplemental Figure 2

#### Supplemental Figure 2

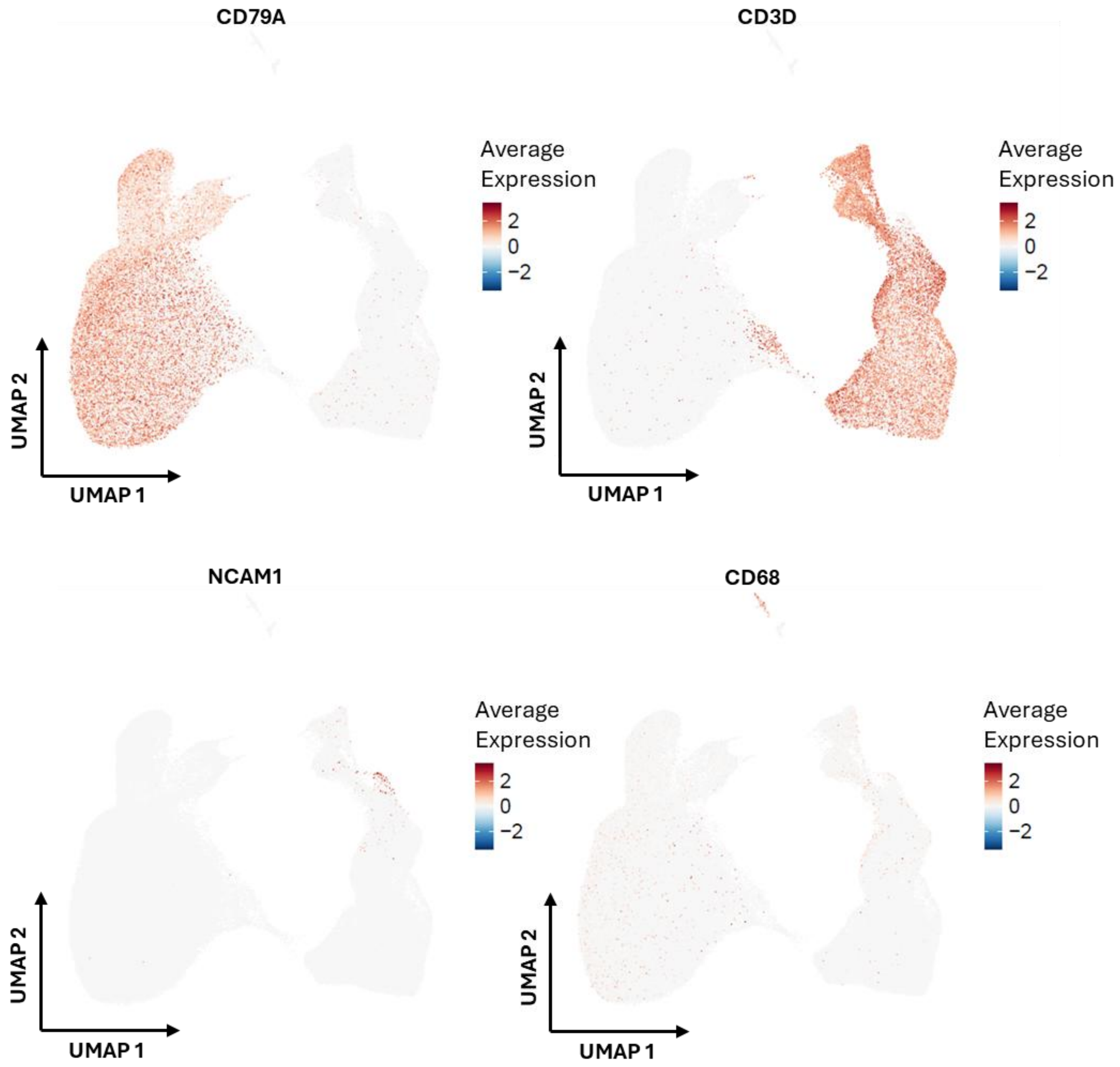

### Supplemental Figure 3

#### Supplemental Figure 3

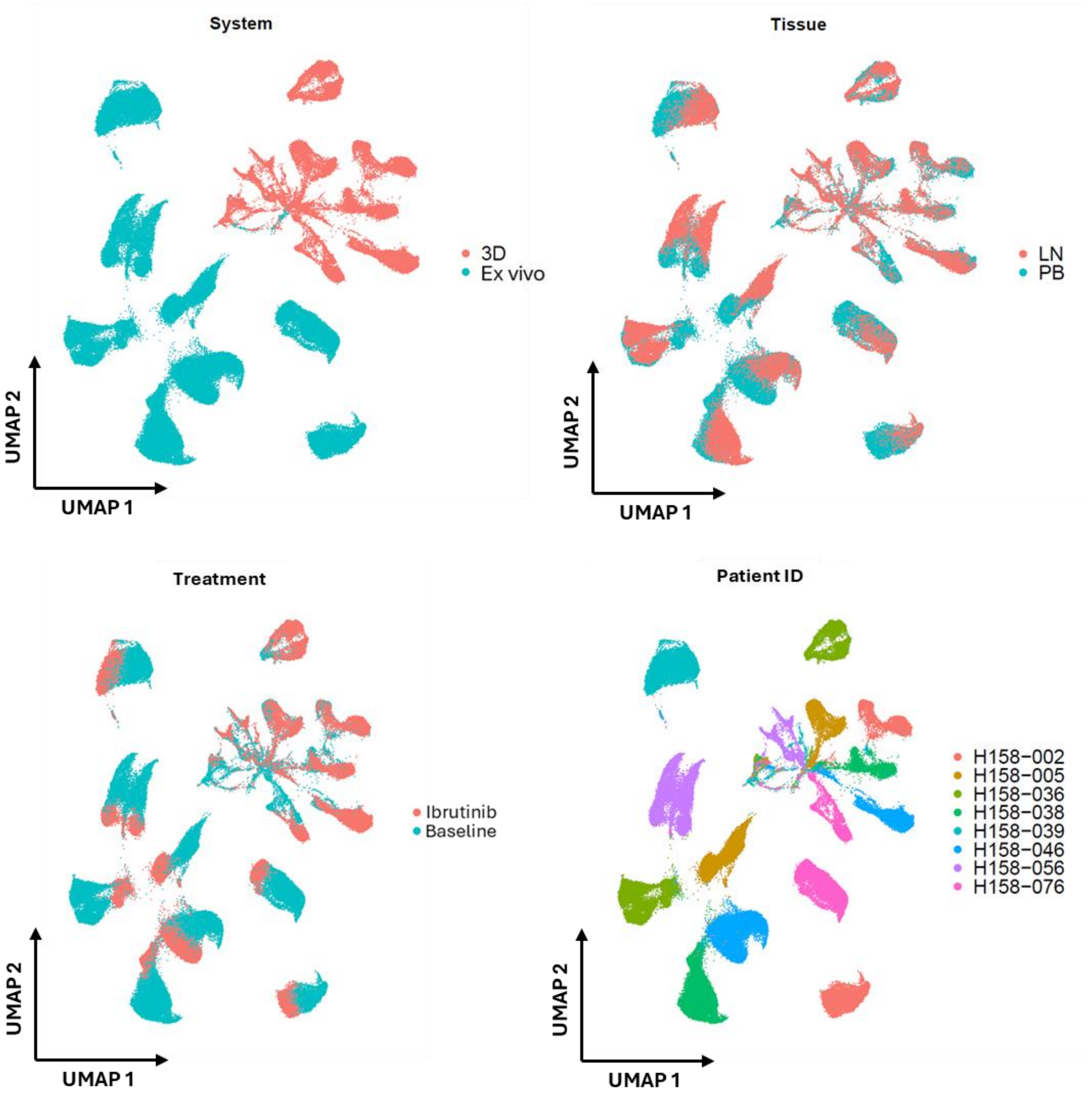

### Supplemental Figure 4

#### Supplemental Figure 4

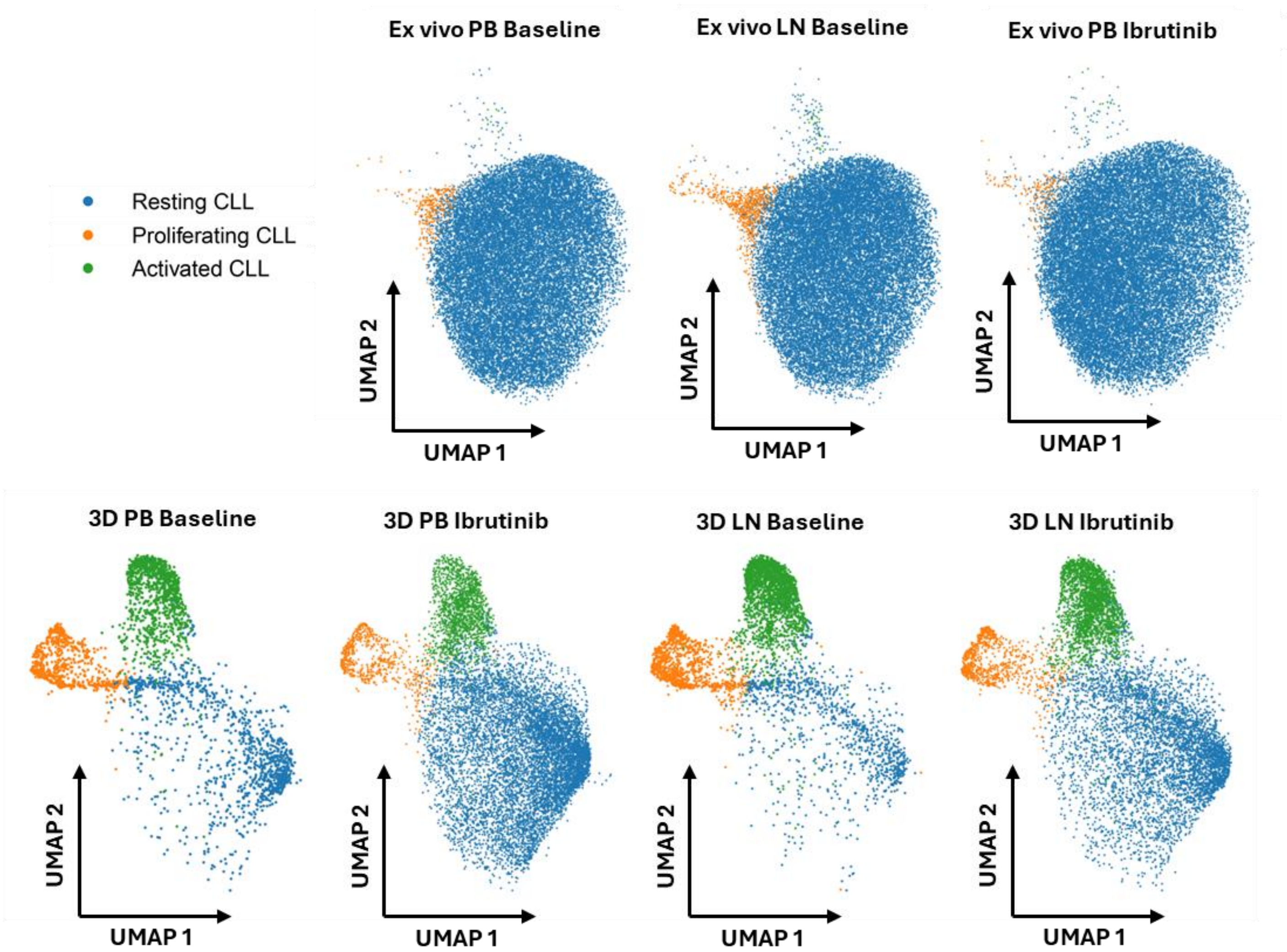

### Supplemental Figure 5

#### Supplemental Figure 5

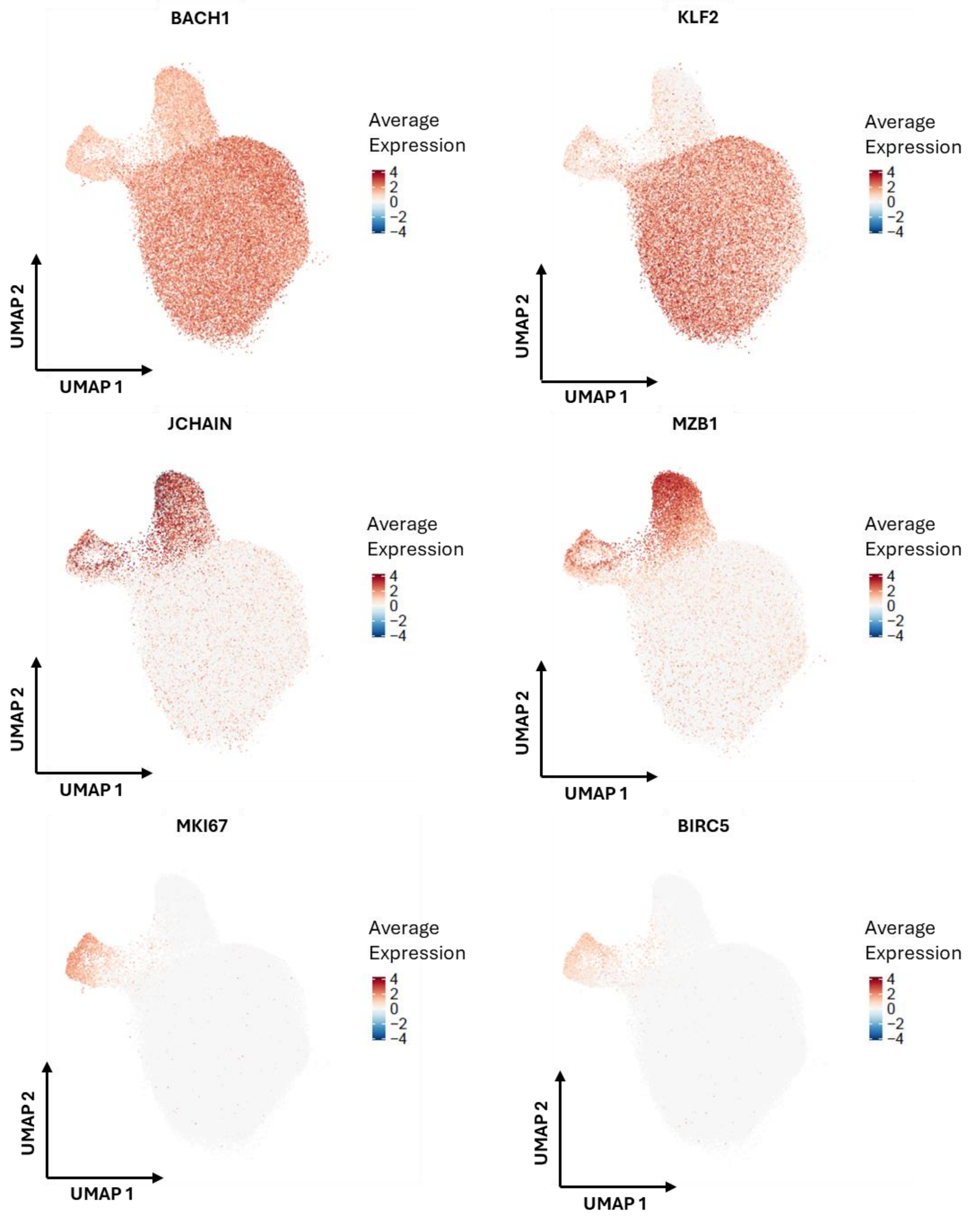

### Supplemental Figure 6

#### Supplemental Figure 6

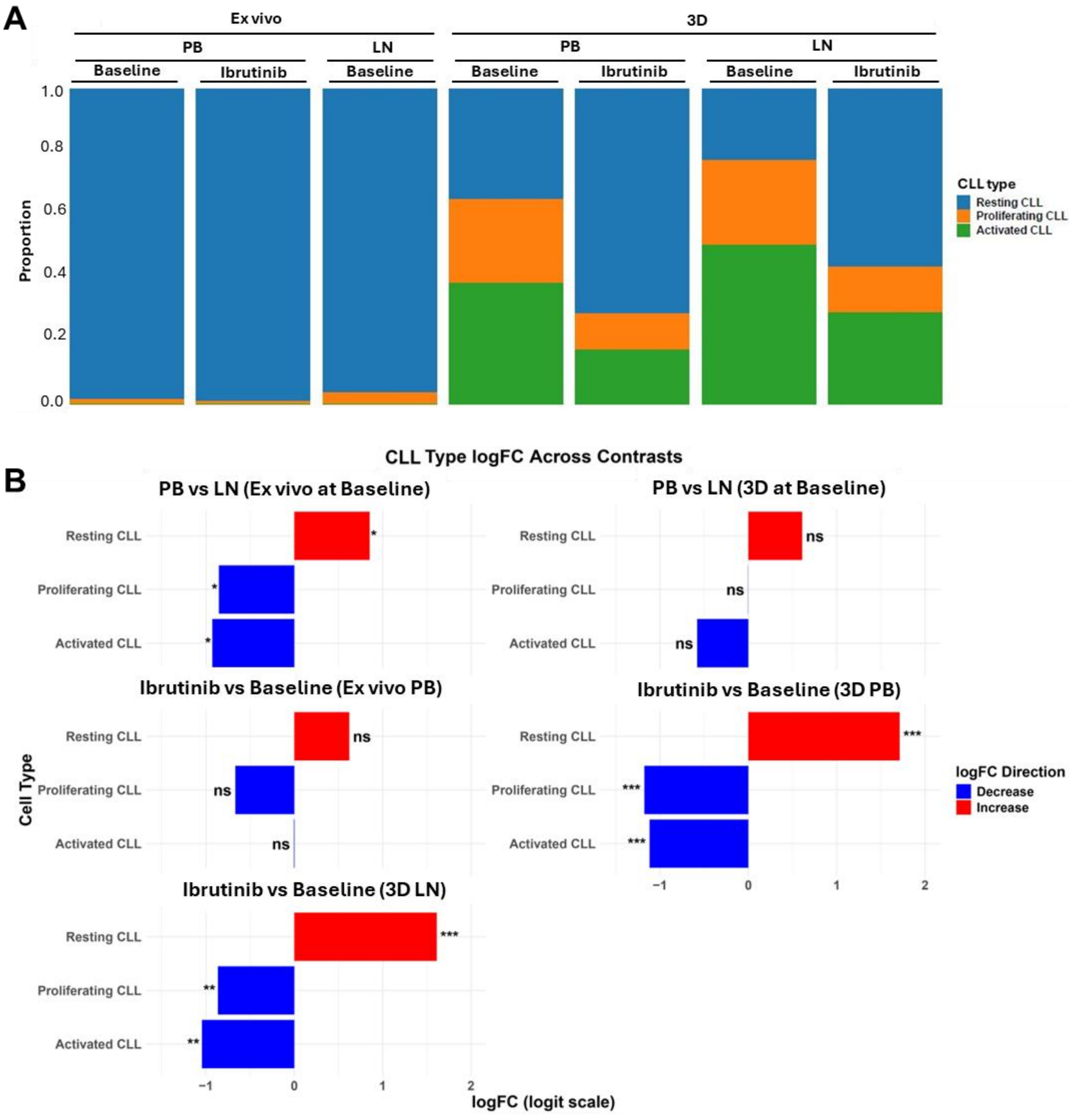

### Supplemental Figure 7

#### Supplemental Figure 7

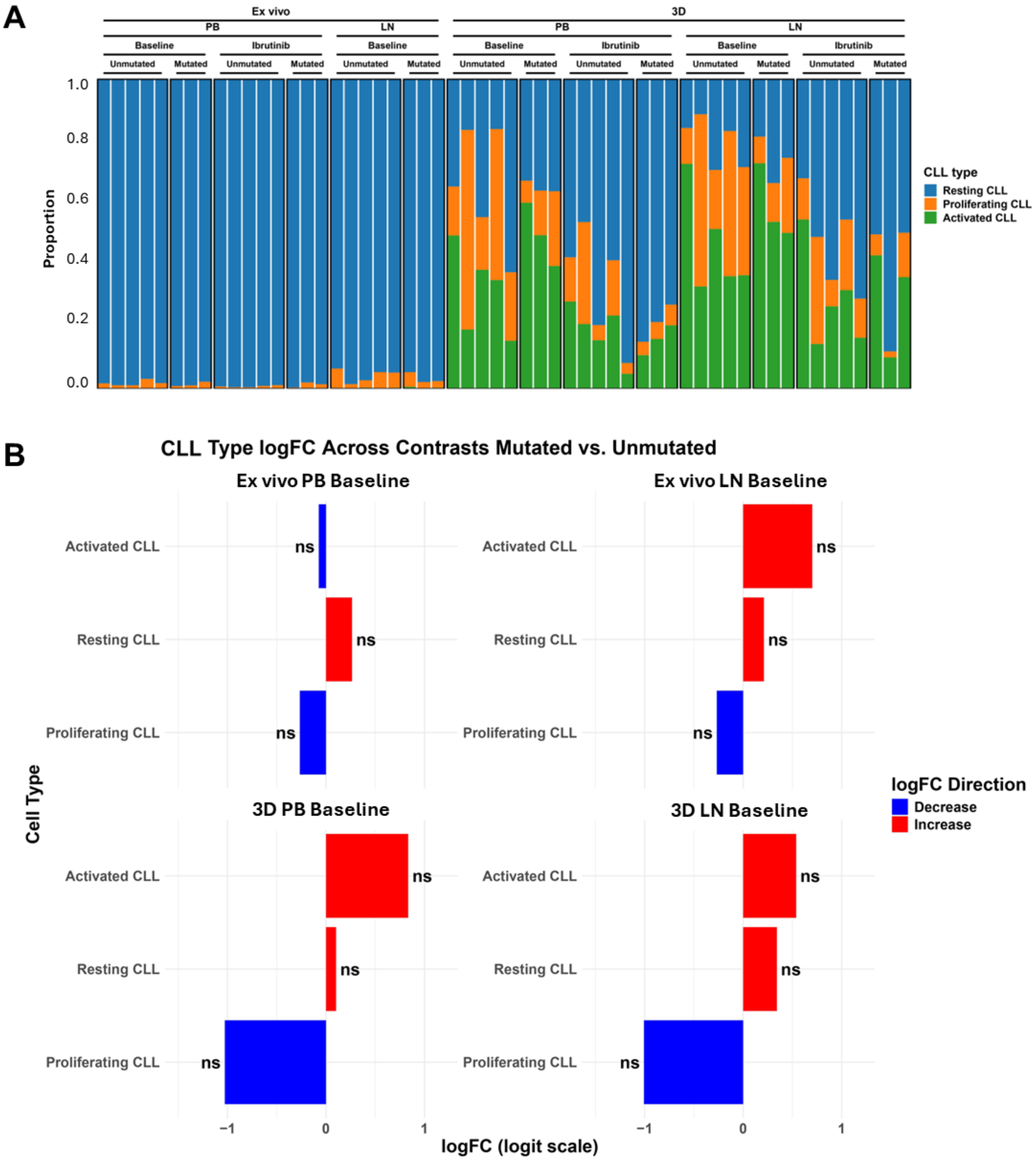

### Supplemental Figure 8

##### Supplemental Figure 8

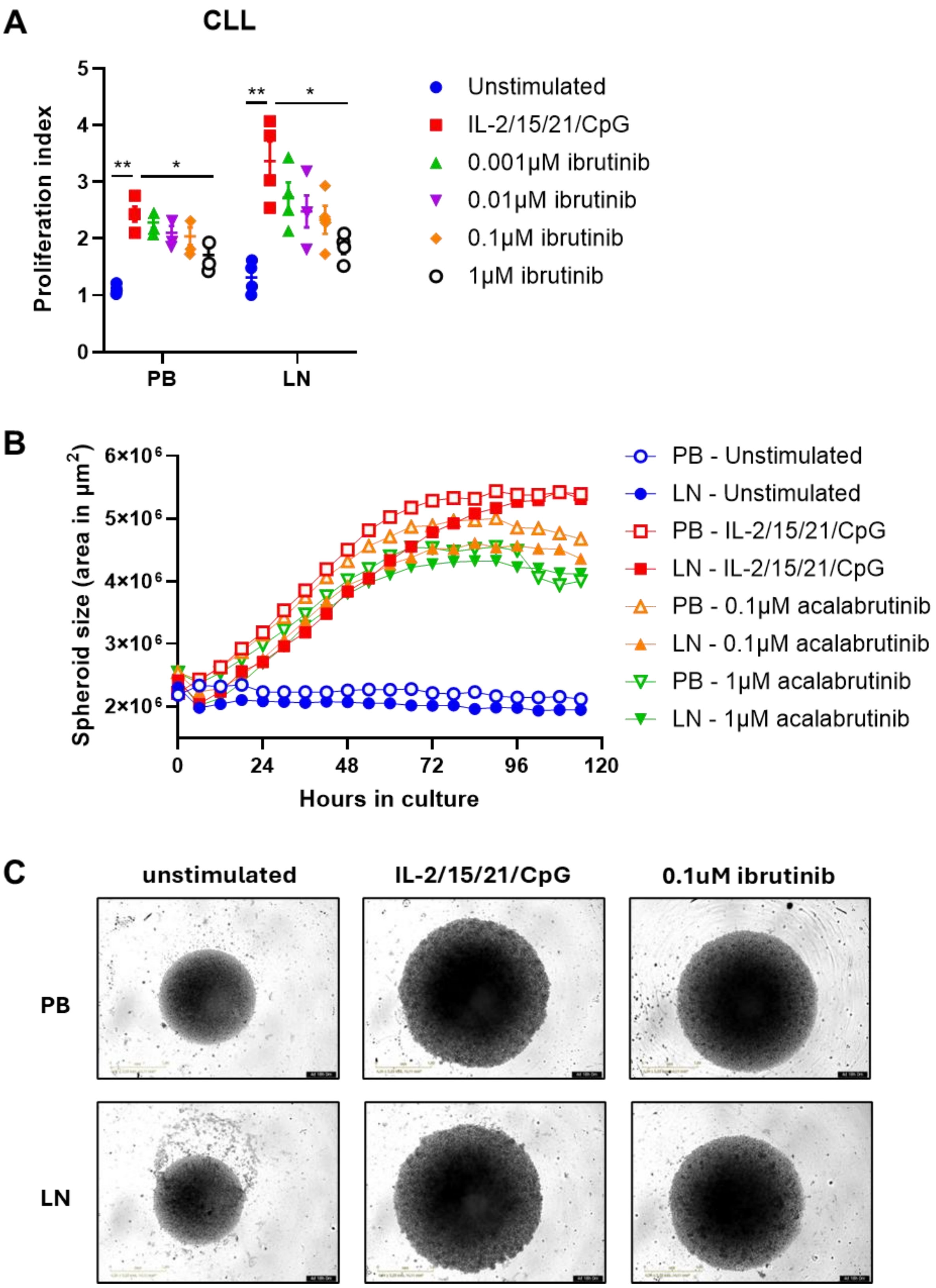

### Supplemental Figure 9

#### Supplemental Figure 9

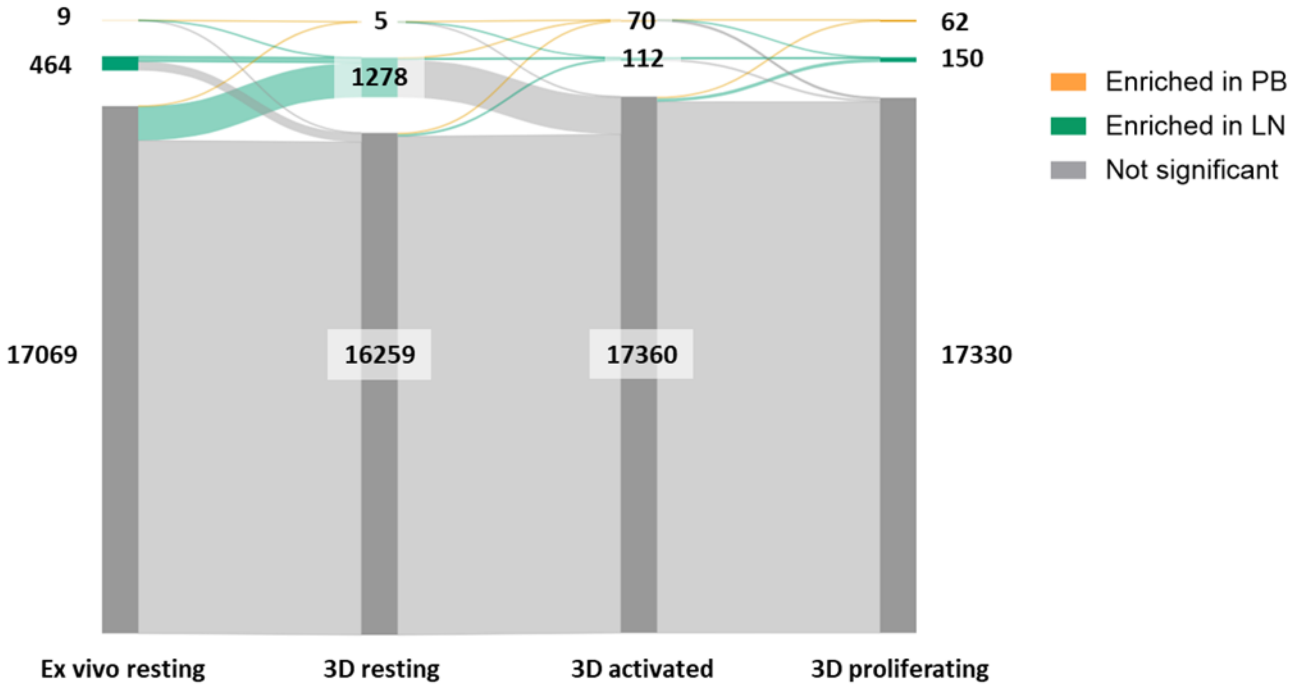

### Supplemental Figure 10

#### Supplemental Figure 10

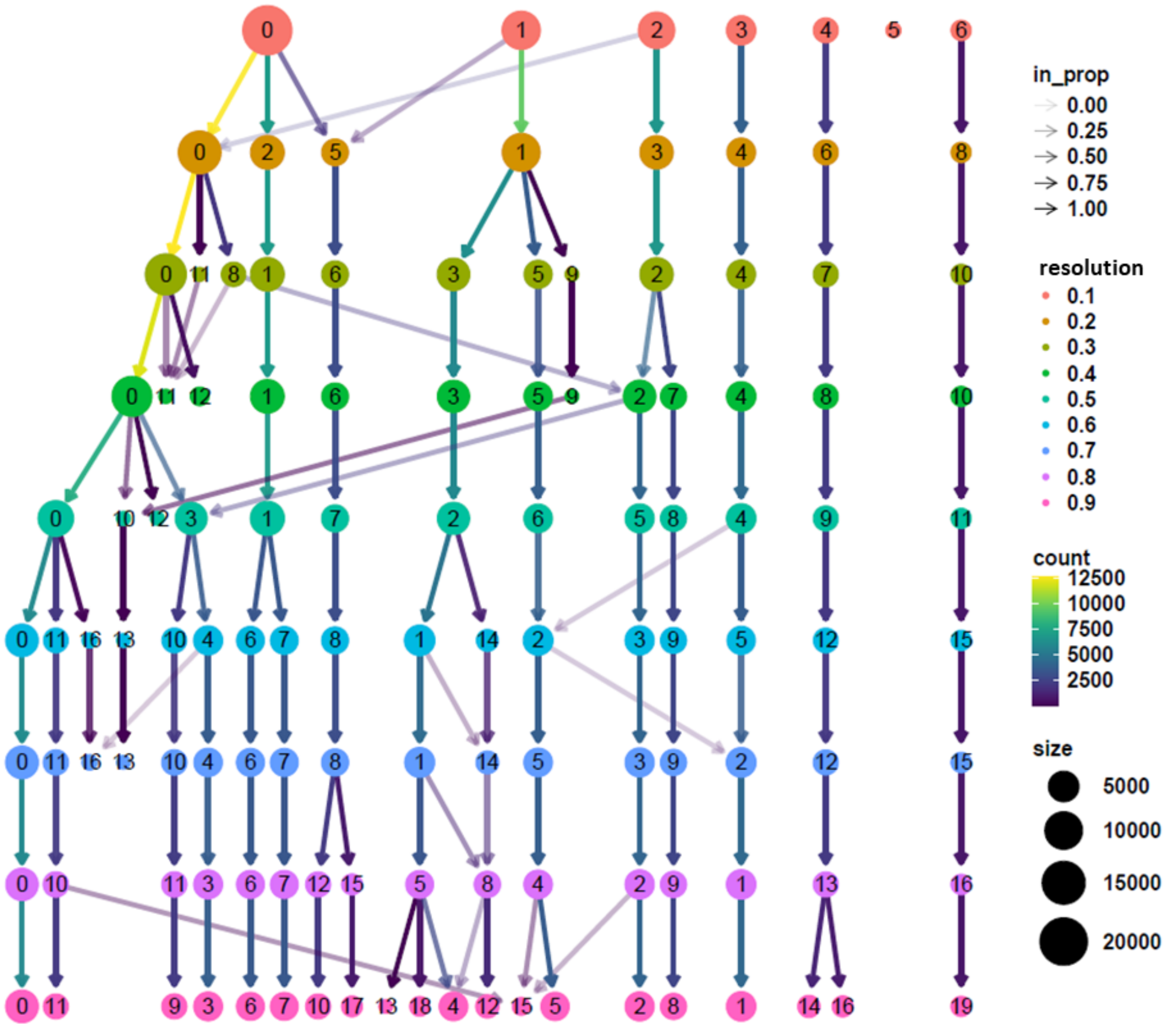

### Supplemental Figure 11

#### Supplemental Figure 11

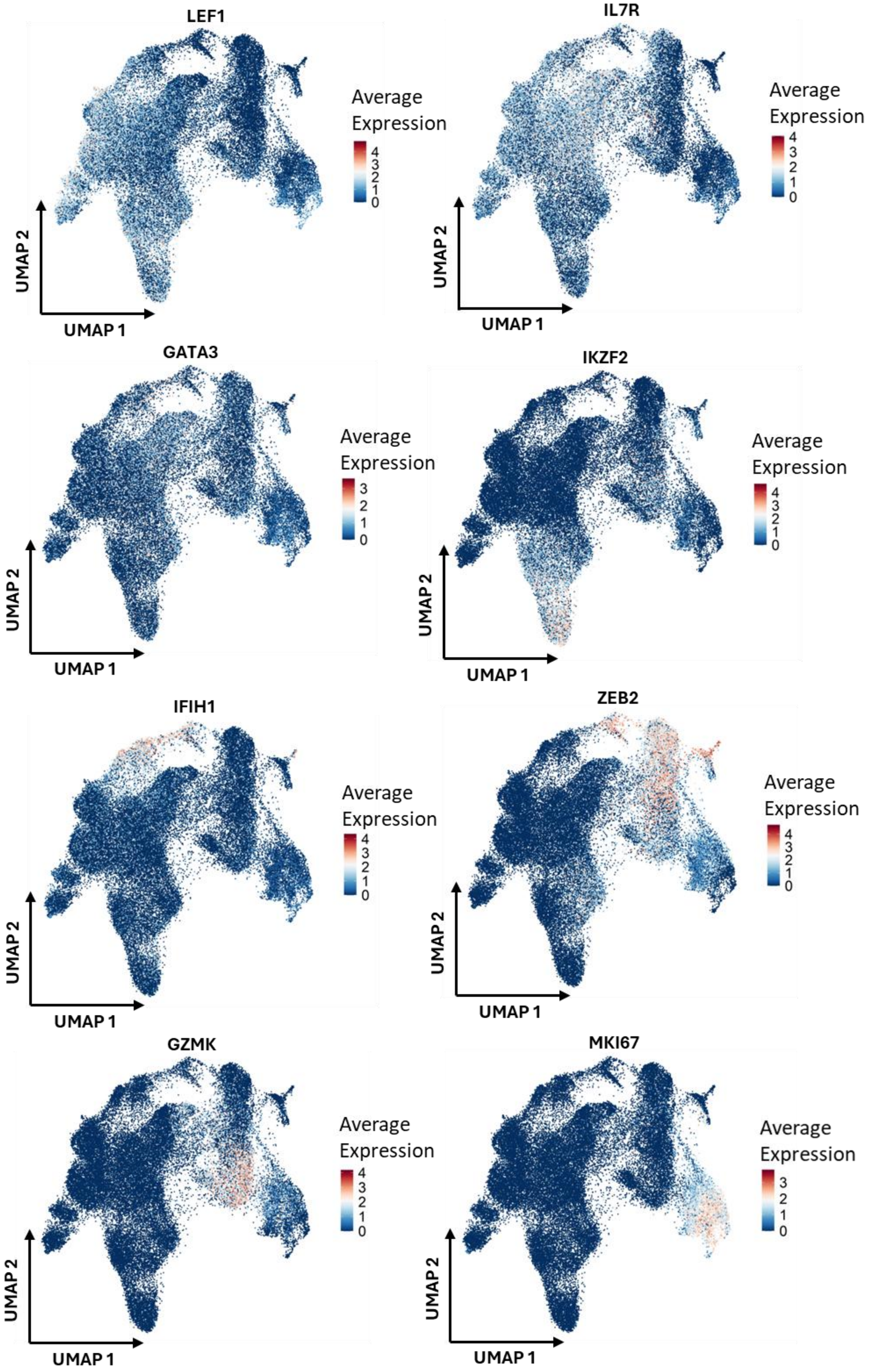

### Supplemental Figure 12

#### Supplemental Figure 12

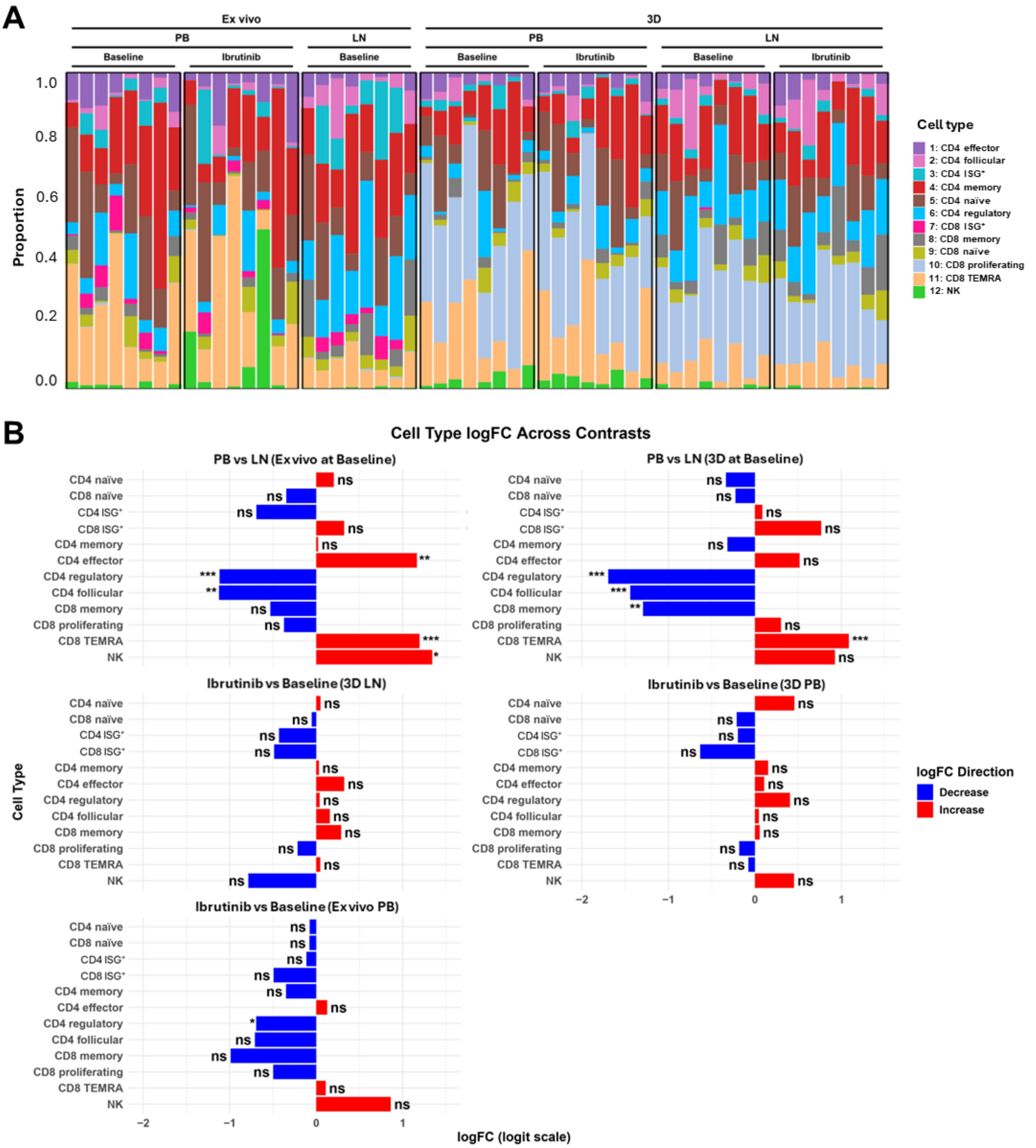

### Supplemental Figure 13

#### Supplemental Figure 13

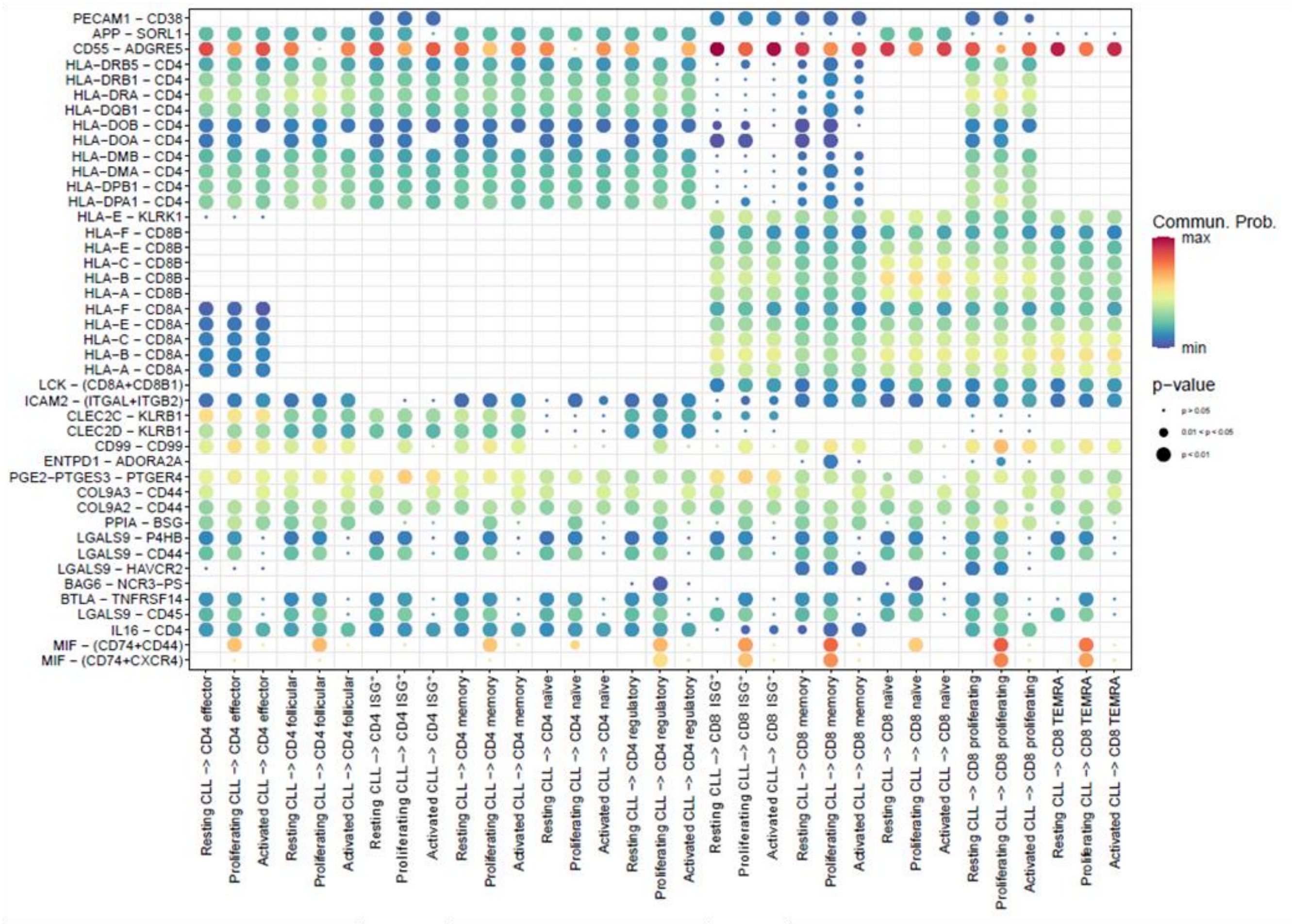

### Supplemental Figure 14

#### Supplemental Figure 14

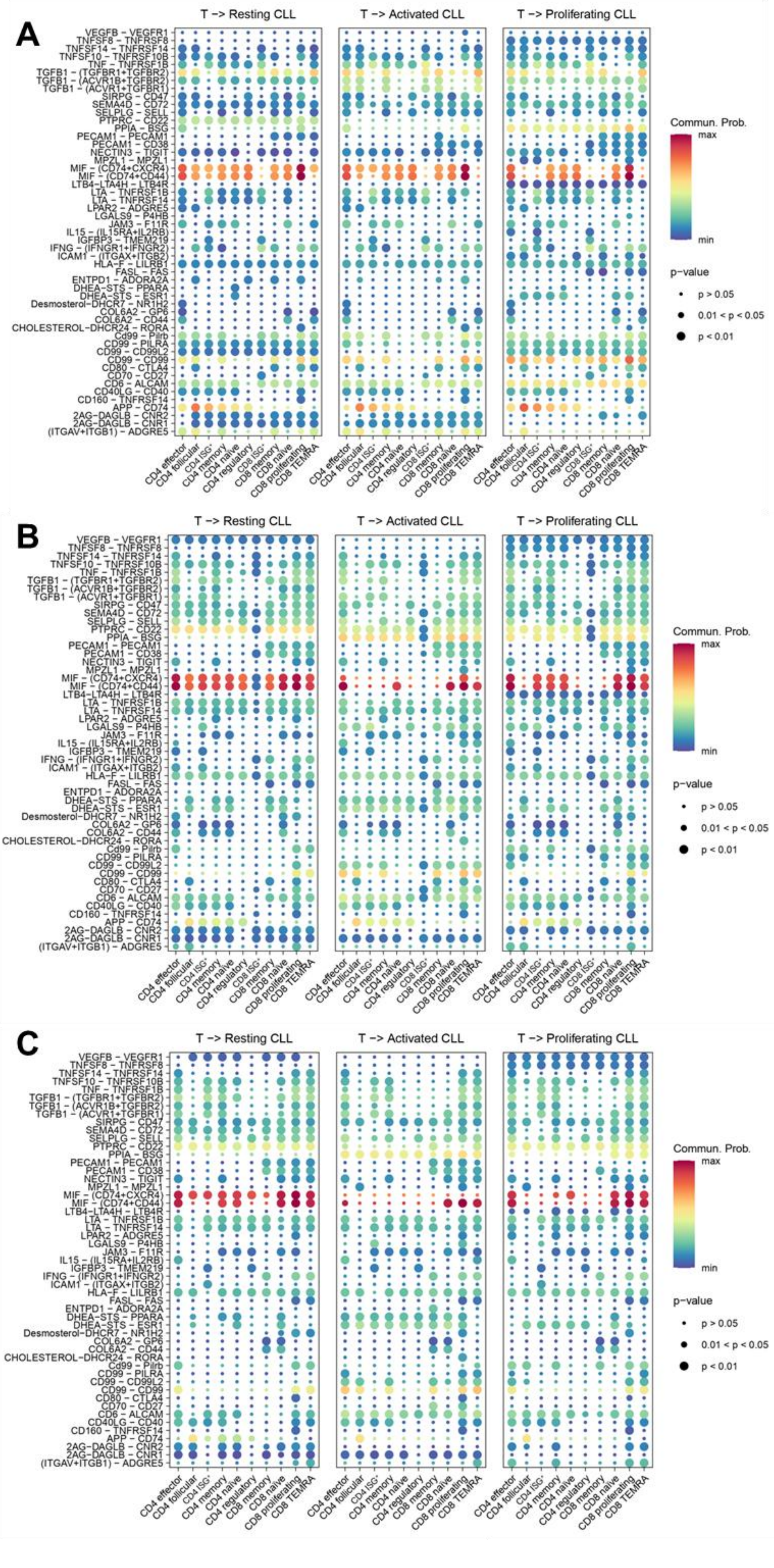

### Supplemental Figure 15

#### Supplemental Figure 15

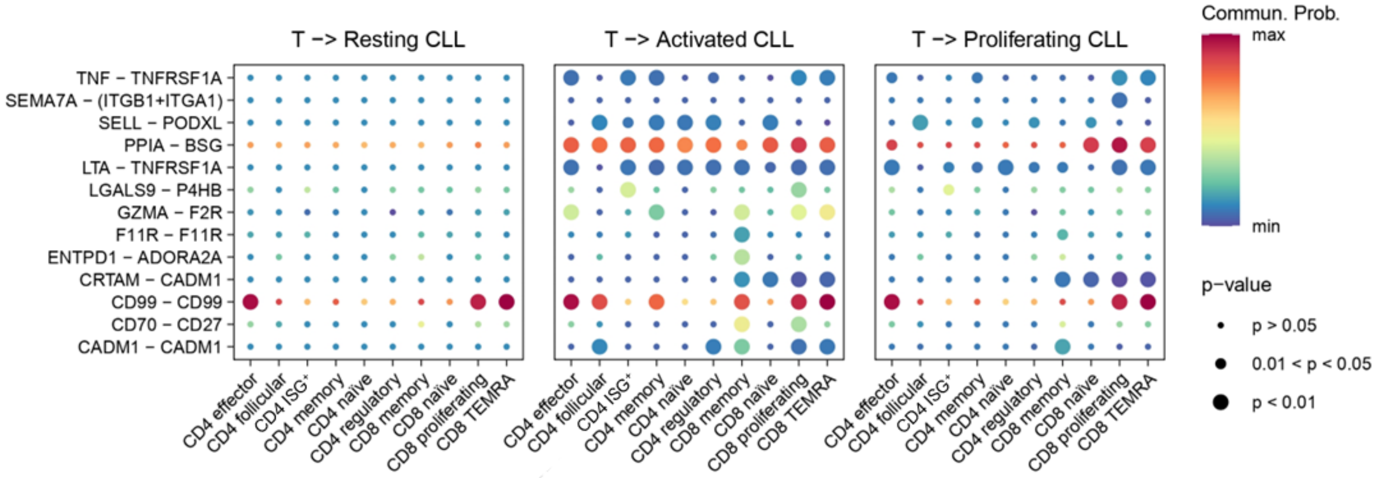

### Supplemental Figure 16

#### Supplemental Figure 16

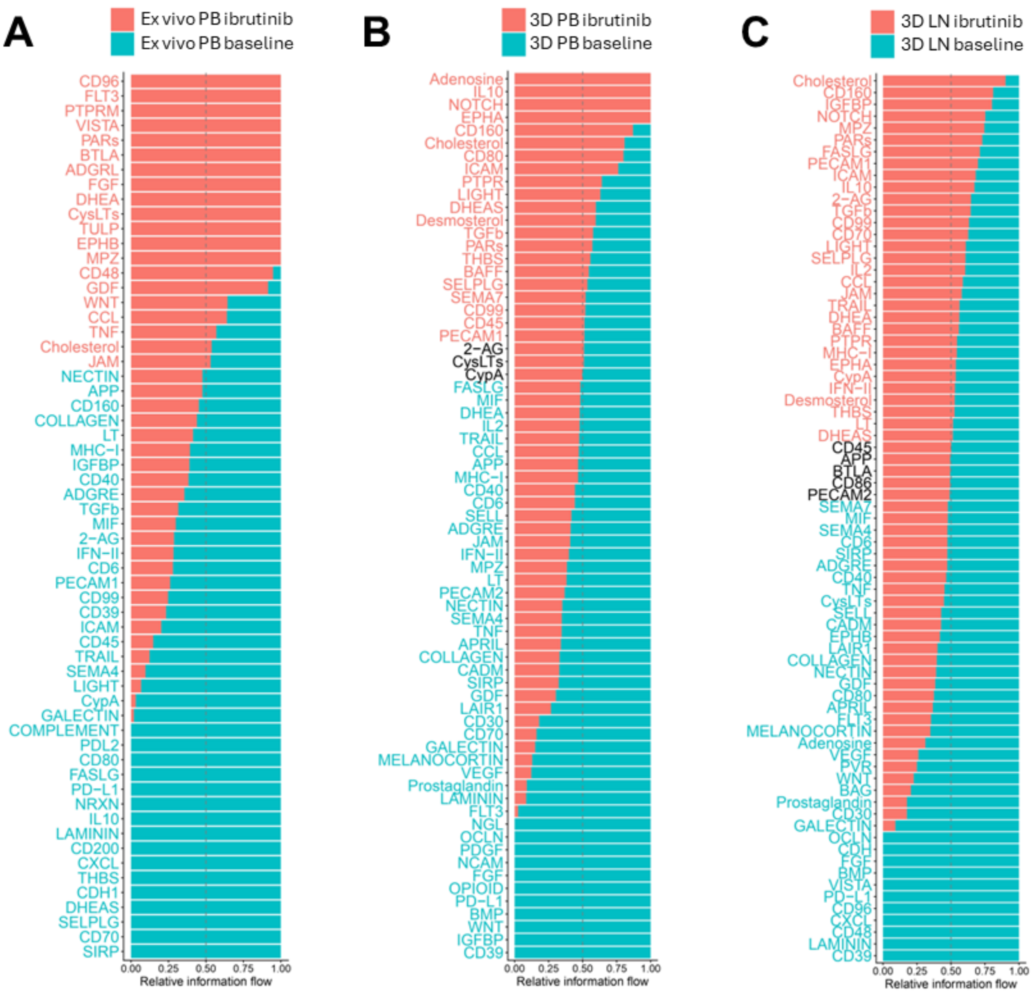

### Supplemental Figure 17

#### Supplemental Figure 17

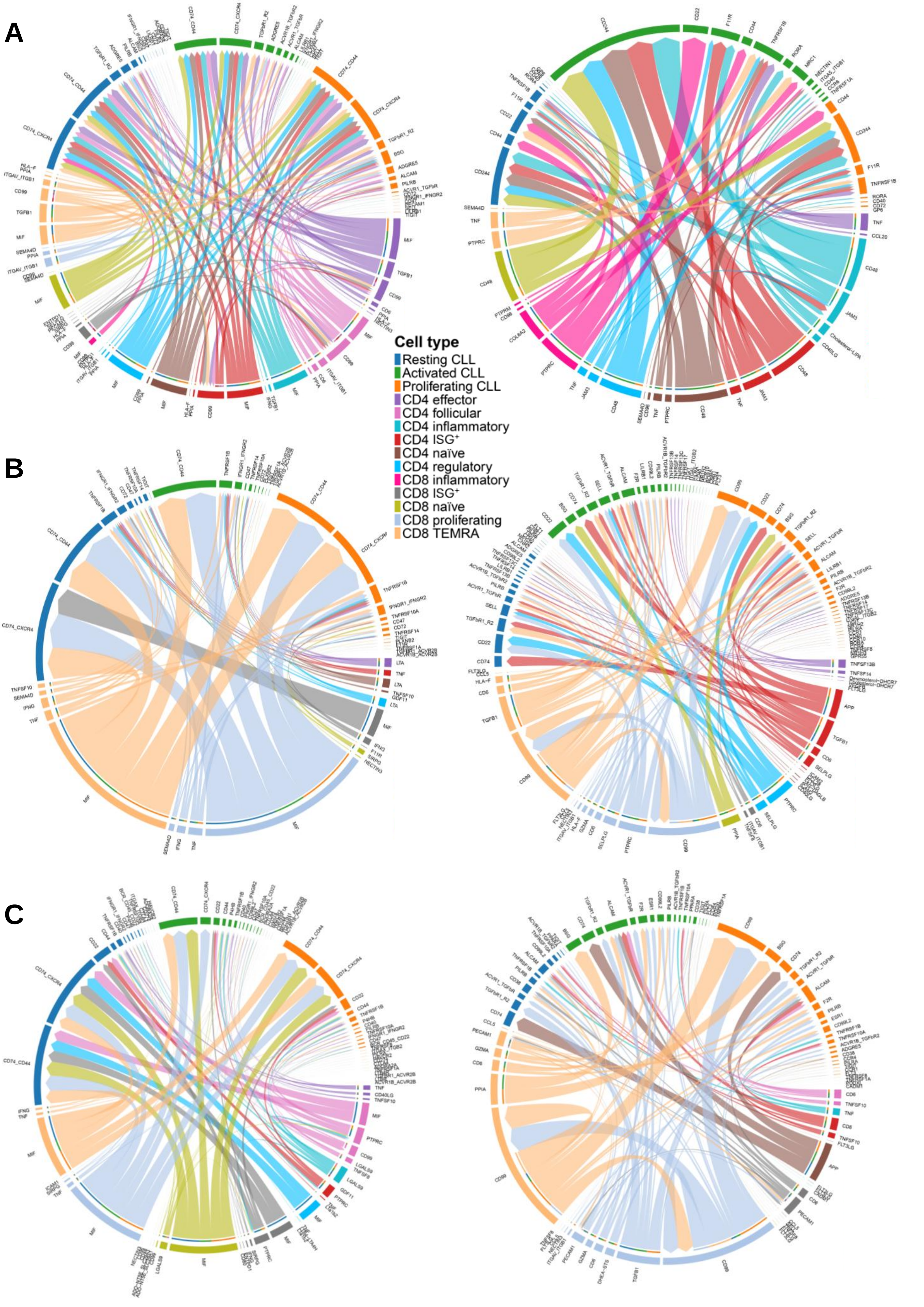

### Supplemental Figure 18

#### Supplemental Figure 18

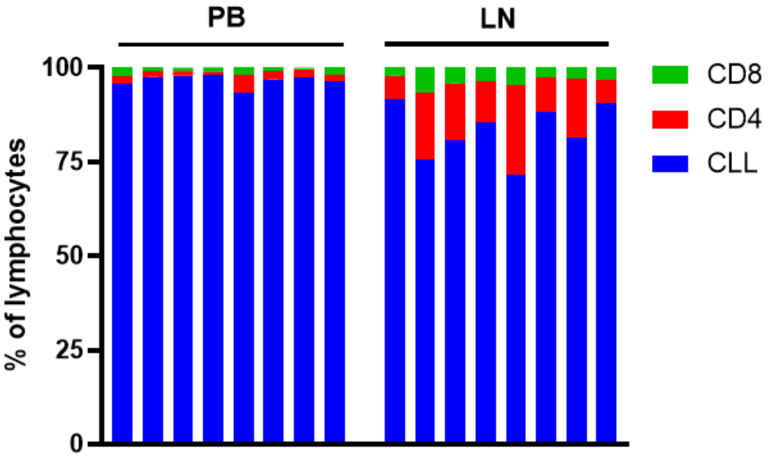
